# Cooperativity among clustered κB sites within promoters and enhancers dictates transcriptional specificity of NF-κB RelA along with specific cofactors

**DOI:** 10.1101/2024.07.03.601930

**Authors:** Shandy Shahabi, Tapan Biswas, Yuting Shen, Rose Sanahmadi, Yaya Zou, Gourisankar Ghosh

## Abstract

Non-consensus binding sites of transcription factors are often observed within the regulatory elements of genes; however, their effect on transcriptional strength is unclear. Within the promoters and enhancers of NF-κB-responsive genes, we identified clusters of non-consensus κB DNA sites, many exhibiting low affinity for NF-κB in vitro. Deletion of these sites demonstrated their collective critical role in transcription. We explored how these “weak” κB sites exert their influence, especially given the typically low nuclear concentrations of NF-κB. Using proteomics approaches, we identified additional nuclear factors, including other DNA-binding TFs, that could interact with κB site-bound NF-κB RelA. ChIP-seq and RNA-seq analyses suggest that these accessory TFs, referred to as the TF-cofactors of NF-κB, facilitate dynamic recruitment of NF-κB to the clustered weak κB sites. Overall, the occupancy of NF-κB at promoters and enhancers appears to be defined by a collective contribution from all κB sites, both weak and strong, in association with specific cofactors. This congregation of multiple factors within dynamic transcriptional complexes is likely a common feature of transcriptional programs.

**SIGNIFICANCE:** The NF-κB RelA dimers undergo rapid activation by cytokines and pathogens, driving expeditious expression of target genes upon binding to DNA elements known as κB sites, located in the regulatory regions. We find that promoter and enhancer regions of RelA target genes harbor multiple κB sites, most being non-consensus with minimal affinity to NF-κB in vitro. Recruitment of RelA dimer in vivo depend on these κB sites, weak and strong, and appears to be regulated by various accessory factors, including other DNA-binding transcription factors. Overall, this study points to a coordinated network of factors communicating with both weak and strong κB sites to recruit RelA dimers, enabling rapid gene activation.

## INTRODUCTION

Transcription is triggered by transcription factors (TFs) binding to specific short DNA response elements (REs) located commonly within the non-coding promoter and enhancer regions (1,2). Our understanding of interactions between eukaryotic TFs and their cognate REs in genes have improved significantly due to genome-wide and imaging analyses of the last two decades (3-5). These studies have also underscored various unresolved aspects about sequence-specific DNA recognition by transcription factors in cells (5-7). The first intriguing aspect is the presence of multiple TF-binding sites of a wide range of affinities in a promoter, which raises the question of the possible roles of weak binding sites in transcription. The second intriguing aspect is the association of many TFs to promoter and enhancer regions that lack cognate DNA binding sites of these TFs, hinting at their non-DNA binding roles in gene expression (8,9). It is unclear if these auxiliary TFs regulate recruitment of primary TF to the weak DNA REs. We addressed these questions through investigation of transcriptional programs mediated by NF-κB RelA, a suitable model system due to availability of information from decades of rigorous studies.

Among the NF-κB family of dimeric TFs, the p50:RelA heterodimer and the RelA:RelA homodimer are most ubiquitous and well-studied (10,11). These dimers bind with high affinity to ∼10-bp sequences (κB sites), with a general consensus of GGGRN<u>N</u>YYCC, defined over 35 years ago (12,13). Several studies indicate binding of various NF-κB dimers to thousands of degenerate κB sites with a wide gradient in affinity (10,14-16). Some the weak sites are not easily recognizable nonetheless bearing specificity to specific NF-κB dimers. The promoters of known NF-κB target genes often contained several of these weak binding sites; however, the functional importance of the vast majority of these weak sites remains unexplored. The promoters of prominent NF-κB regulated genes e.g., E-selectin, Cxcl10, Ccl2, IL-1β, TNF-α, and Nfkbia, all contain multiple κB sites including many of lower affinity. Cell-based reporter assays indicated the necessity of some of these weak sites in optimal reporter expression (10,17). However, the prevalence of bona fide weak κB sites in native promoters and enhancers of the above-mentioned and other NF-κB-regulated genes is largely unclear, and the lower limit of binding affinity of a weak κB site for it to be functional is not defined. Essential roles of weak DNA binding sites of various transcription factors within the enhancer regions have been reported during development (18-22), where suboptimal binding of TFs to the weak sites appears to provide specificity for gene expression. For the TF ETS, a single nucleotide change in a binding site causing only a small enhancement in the binding affinity is observed to cause pathogenicity (22). However, the mechanistic possibilities of how multiple binding sites within a promoter/enhancer can act in concert, either for the strength or the specificity of transcription, remains largely unexplored.

TFs have also been observed to require auxiliary factors for cognate DNA recognition (23). Previous studies revealed that NF-κB is recruited along with additional factors (10,24-26). The associated factors during RelA recruitment are promoter and cell type specific although it is unclear if they play a role in facilitating occupancy (24,25,27).

We set out to clarify these unresolved issues in transcription through investigating the role of weak κB sites and supporting factors in the recruitment of NF-κB RelA to DNA, and consequently transcription. Through analysis of ChIP-seq data of NF-κB p50:RelA and RelA:RelA dimers during stimulus-dependent expression of NF-κB-responsive genes, we observed that multiple κB sites are present in clusters within ∼150-bp long DNA segments of promoter and enhancer regions. Additionally, we observed many of these κB sites to be weak i.e., displaying a barely detectable affinity in vitro. However, these sites appear to play essential roles in recruiting NF-κB to target genes and driving transcription. Additionally, we observe that numerous protein factors, including other TFs (TF-cofactors), RNA-binding proteins, and enzymes associate with NF-κB RelA on DNA. Further investigation of these TF-cofactors revealed their critical role in enhancer- and promoter-specific recruitment of RelA.

## RESULTS

### Clusters of κB sites of a wide range in affinity are present in promoters of NF-κB target genes and these collectively dictate recruitment of RelA to DNA

We examined the relationship between ChIP-seq scores of RelA, reflecting occupancy of RelA on DNA in the cell, and sequence of κB sites within the promoters of over 100 NF-κB-regulated murine genes that are stimulated by TNF-α within a 30-minute stimulation period, as reported by Ngo et al (28). The sites are designated as κB sites if they matched any of the 1800 sites with a measurable Z-score derived from an earlier protein microarray-based assay (PBM) analysis (14). We characterized the strong and weak κB sites of the promoter and enhancer regions to assess how a set of these κB sites could facilitate transcription through recruitment of RelA dimer in association with accessory cofactors. The κB sites were designated as strong or weak considering both their physical and functional characteristics. Binding characteristics of A-centric (IFNβ-κB) and G-centric (STX11-κB) κB sites towards p50:RelA heterodimer or RelA:RelA homodimer were used as benchmarks (13). In general, κB-site with affinity (K_D_) to NF-κB RelA dimers above and below 100 nM - assessed in vitro using a solution-based binding assay at near-physiological ionic strength and pH - were defined as weak and strong, respectively. The Z score of binding of 1800 κB sites to NF-κB RelA dimer estimated in the PBM assay (14) were tallied to it, and an arbitrary Z score threshold of ∼ 6.0 separated weak and strong κB sites. Additionally, a transcriptional activation potential of a single κB site in luciferase reporter-based assay to less or more than 2-fold were considered as functionally weak and strong, respectively. The spectrum of considered κB sites deviated from the 10-bp consensus at the most at three positions. Strikingly, we observed that not all strong κB sites are aligned with RelA ChIP peaks. Rather, we observed that the 500-bp window centered around the ChIP-seq peak for most TNF-α-responsive genes contained a strong κB site surrounded by multiple weak κB sites (**Figure 1A**). Analysis of the weak κB site abundance around RelA ChIP-seq peaks sorted by intensity (ranked as top and bottom) revealed a significantly higher number of weak κB sites around top ranked RelA peaks (**Figure 1B**). However, we observed only a weak correlation between RelA ChIP-seq score and the Z-score of the strong κB site proximal to the central peak motif (**Supplementary Figure 1A**). This suggested the possibility that individual affinity of the central strong κB site is insufficient in predicting occupancy of RelA dimers to promoter regions in cells. Intriguingly, we observed a stronger correlation of ChIP-seq strength with calculated combined Z-score of sites in an extended window of up to 500 bp around the central strong κB motif (**Supplementary Figure 1A**). The correlation reduced with exclusion of the contribution from weak κB motifs, suggesting a significant contribution of the colocalized weak κB motifs in recruitment of RelA (**Figure 1C**). The clusters of κB sites primarily spanned about 150 to 200 bp within the analyzed window of 500 bp, thus mimicking the span of an open chromatin observed within promoters and enhancers during transcription (29).

**Figure 1.**
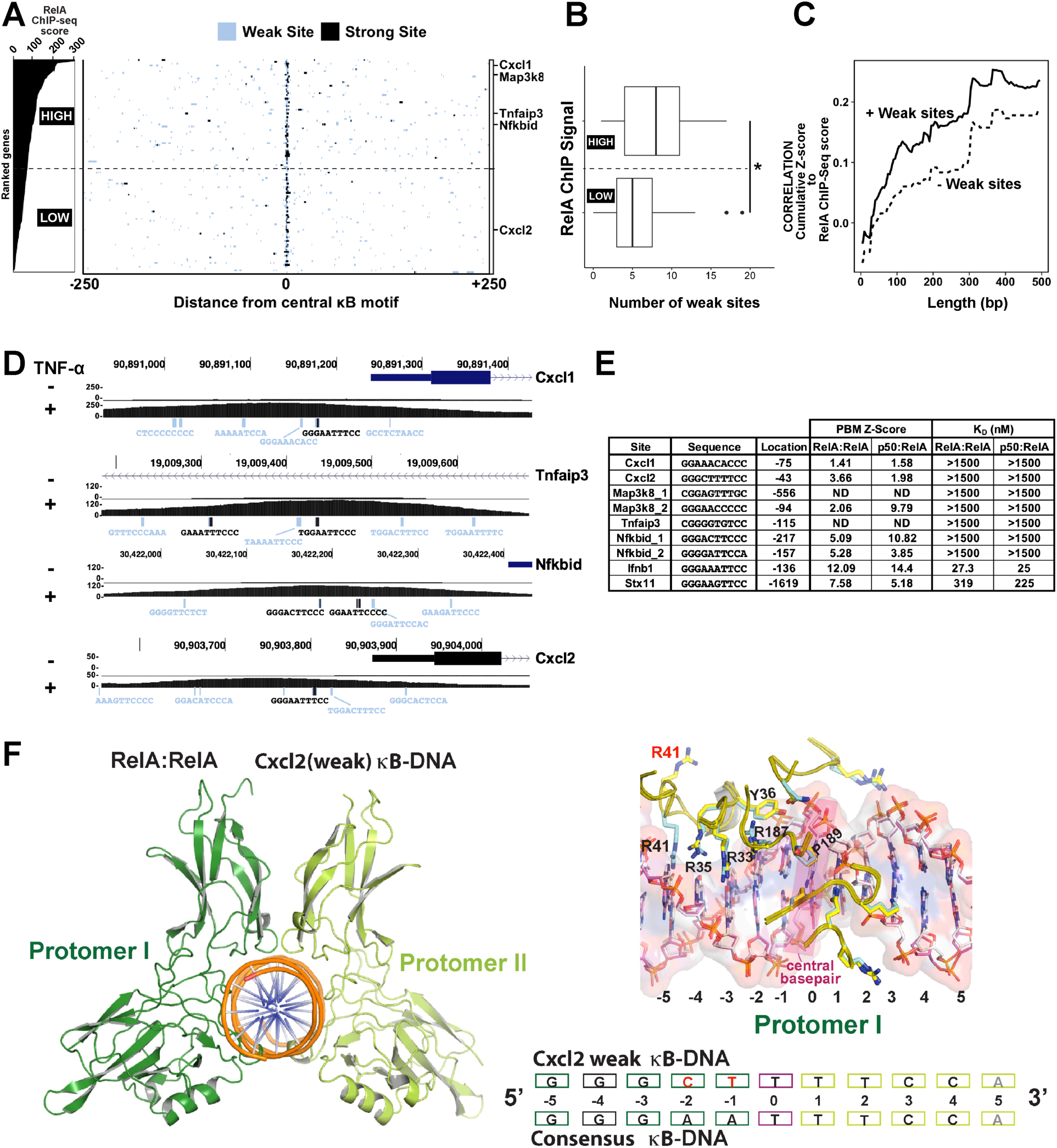
Identification and analysis of weak κB motifs in RelA-binding promoter sites of TNF-α-responsive genes. **A.** Left panel, Genes ranked based on RelA ChIP-seq scores in the analyzed promoter regions (separated by a dotted horizontal line in two halves for high and low). Right panel, 500-bp regions centered around the strongest RelA motif within RelA ChIP-seq peaks (Ngo, et al. 2020) were analyzed for the presence of strong (black) and weak (blue) κB motifs, as defined by a cut-off Z-score of 6 in PBM analysis of p50:RelA heterodimer (Siggers, et al. 2009). n =116 RelA peaks analyzed. **B.** Boxplot of number of weak κB sites in the top half versus bottom half of genes ranked by ChIP-Seq score. *P < 0.05. P value was calculated by two-tailed Student’s t-test. **C.** Dependence of the “Pearson’s correlation between RelA ChIP-seq score and cumulative promoter Z-score surrounding the central κB site” to “analysis window length” of 10-500 bp. Analysis either included (solid) or excluded (dashed) contribution of weak κB motifs. **D.** Genome browser tracks showing κB motifs and their sequences within a 500 bp window of Cxcl1, Cxcl2, Tnfaip3, and Nfkbid promoters, along with TNF-α induced RelA ChIP-seq signal. Strong and weak κB site sequences in black and blue, respectively. RelA ChIP-seq signal intensity is indicated in the left. **E.** BioLayer interferometry assay derived equilibrium binding affinity (primarily from this study) with Z-scores (Siggers, et al. 2009) of κB sites for RelA homodimer and p50:RelA heterodimer. **F.** A cartoon representation of the RelA homodimer (left) bound to a weak κB site derived from Cxcl2 gene (bottom). Structural similarity/dissimilarities in interaction of the RelA protomers to weak versus strong κB motif (right).

We further analyzed the arrangement of κB sites in promoters of prominent NF-κB-responsive murine genes Cxcl1, Cxcl2, Nfkbid, and Tnfaip3 (**Figure 1D**). Both Cxcl1 and Cxcl2 contain an identical strong site (GGGAAATTCC) but the overall ChIP score at the Cxcl2 promoter is noticeably lower than that at Cxcl1 (250 vs 50). This may be reflective of a higher number of weak κB sites in the Cxcl1 promoter compared to Cxcl2 promoter (13 vs 8). The composite cumulative Z-score of Cxcl1 weak sites for RelA dimers is also higher than that of Cxcl2 (70.6 vs 58.5). Some of these weak sites, such as GGA<u>T</u>TTT<u>TT</u> and GGGAATTT<u>TA</u> (deviations from consensus are underlined) found near the strong site in Cxcl1 and Tnfaip3 promoters, displayed minimal resemblance to the κB consensus. To assess if these putative weak sites engage to RelA dimers, we measured affinities of selected κB sites for both NF-κB RelA homodimer and p50:RelA heterodimer (of wild-type full-length proteins) at a near-physiological salt concentration by biolayer interferometry (BLI) assays. Both NF-κB dimers bound these putative sites with a much-reduced affinity when compared with binding to IFN-β1: GGGAAATTCC (K_D_ ∼ 27 and Z-score ∼14), and STX11: GGGAAGTTCC (K_D_ ∼250 and Z-score 5.8) sites as references of strong and weak κB sites. Although minor binding was detected for most weak sites at higher protein concentrations, precise K_D_ values could not be determined. Two of the sites tested did not display any binding even at the highest protein concentration (**Figure 1E** and **Supplementary Figure 1B**) and these are estimated to have K_D_ >1.5 μM. Since the nuclear concentration of RelA upon TNF-α stimulation is observed to be around 200-250 nM (**Supplementary Figure 1C**), only about 6% of these dimers are projected to bind if K_D_ is around 3.0 μM. These findings raised queries on the authenticity and functional relevance of putative weak sites in vivo. It should be noted that there is a discrepancy between binding constants measured with native proteins at near physiological ionic strengths using solution-based methods versus truncated proteins under non-physiological ionic strength using microarray-based methods. We also qualitatively examined the binding strengths of weak and strong sites within the Cxcl2 promoter using electrophoretic mobility shift assay (EMSA) with recombinant p50:RelA heterodimer and RelA:RelA homodimer (**Supplementary Figure 1D**). The results indicated that mutation of both κB sites in Cxcl2 promoter, referred here to as Cxcl<u>2 d</u>ouble <u>m</u>utant (2DM), abrogated binding of NF-κB completely, while mutation of only the weak site (2ΔW) did not cause any discernible effect. Sequence-specific binding at the weak κB site (2ΔS) was retained upon mutation of strong site for both dimers, demonstrating that NF-κB has the capacity to bind this weak motif. Collectively, these findings led us to conclude that the putative weak sites with detectable binding *in vitro* may have a functional role.

To further evaluate if the binding by NF-κB dimers to weak κB sites is specific and authentic, we determined the crystal structure of RelA homodimer in complex with a weak κB site (^-5^G^-4^G^-3^G^-2^<u>C</u>^-1^<u>T</u>^0^T^+1^T^+2^T^+3^C^+4^C) present in the promoter of the Cxcl2 gene (the non-cognate basepairs are underlined and the pseudo-dyad axis separating the two halves of the κB site is marked ^0^T) (**Supplementary Table 1**). The structure revealed that the binding interaction of RelA dimer to this weak site is remarkably similar to that observed with consensus κB sites (**Figure 1F**; PDB code 9E6W). The Arg33 and Arg35 engaged to the ^-4^G^-3^G of one halfsite, as predicted. However, the presence of non-cognate bases (^-2^C^-1^T compared to ^-2^A^-1^A, the cognate bases for the RelA homodimer) influenced the global DNA architecture such that Arg187 is positioned slightly differently, but more importantly, the R41 could not engage with ^-5^G of this weak site. This altered protein-DNA contact likely entails the weaker interaction, which nonetheless is specific. Taken together, these results reflect an enrichment of weak κB motifs in the regulatory regions of NF-κB target genes, which can indeed be recognized specifically by RelA; however, a more elaborate mechanism must underly an efficient occupancy of these weaker sites.

### Weak sites within κB clusters drive NF-κB-induced transcription

We followed through next by exploring the potential of above-mentioned weak κB sites in activating transcription of NF-κB target genes in cellular milieu. We selectively deleted strong and its neighboring weak κB sites, independently — in the promoters of NF-κB target genes Cxcl1, Cxcl2, Nfkbid, Map3k8, and Tnfaip3 of mouse embryonic fibroblast (MEF) cells using CRISPR-Cas9 genome editing technology (**Figure 2A, 2B**). The placement of weak sites in different genes were different in sequence, orientation, and relative distance from the strong site. Control cells were established by deletion of a random segment away from any κB site, and sequencing of bulk populations indicated the deletions to be averaging around 4-10 bp (**Supplementary Figure 2**). These pooled edited cell lines were grown, and the transcript levels of target genes were measured by quantitative PCR (qPCR) upon treatment with TNFα (10 ng/ml) for 1 h. We observed a significantly reduced level of transcription in all five target genes upon deletion of either the strong or weak sites (**Figure 2B**). To assess if deletion of weak or strong sites in the promoter of one gene could affect the expression of other genes, we analyzed the expression levels of Cxcl2, Map3k8 and Tnfip3 following deletion of κB sites in the Cxcl1 promoter, and analyzed expression of Cxcl1, Cxcl2 and Map3k8 upon deletion of κB sites in the Tnfaip3 promoter. For the most, deletion of κB sites in genes within one chromosome did not alter the expression of genes located on other. However, we observed that deletion of either strong or weak κB site in the promoter of Cxcl1 altered the expression of Cxcl2 (**Supplementary Figure 2B**). These two genes are present in tandem on the same chromosome (chr 4 in humans and chr 5 in mice). Thus, these genes appear to be regulated by the overlapping enhancers, suggesting a potential for local regulatory interactions. Chromatin immunoprecipitation (ChIP) qPCR assays using a specific antibody against RelA with the wild-type and mutated Cxcl1 and Cxcl2 promoters indicated a significant reduction of RelA ChIP signal upon deletion of κB sites, weak or strong, compared to a control site deletion (**Figure 2C**). We are unclear to why mutant weak-site had an equally strong effect as the mutant strong site in case of Tnfaip3. This further indicates that weak sites are not always redundant, and one possibility is that this weak site strongly engages another factor critical to RelA recruitment. These results support a critical function of weak sites in recruitment of NF-κB RelA to promoter and in transcription.

**Figure 2.**
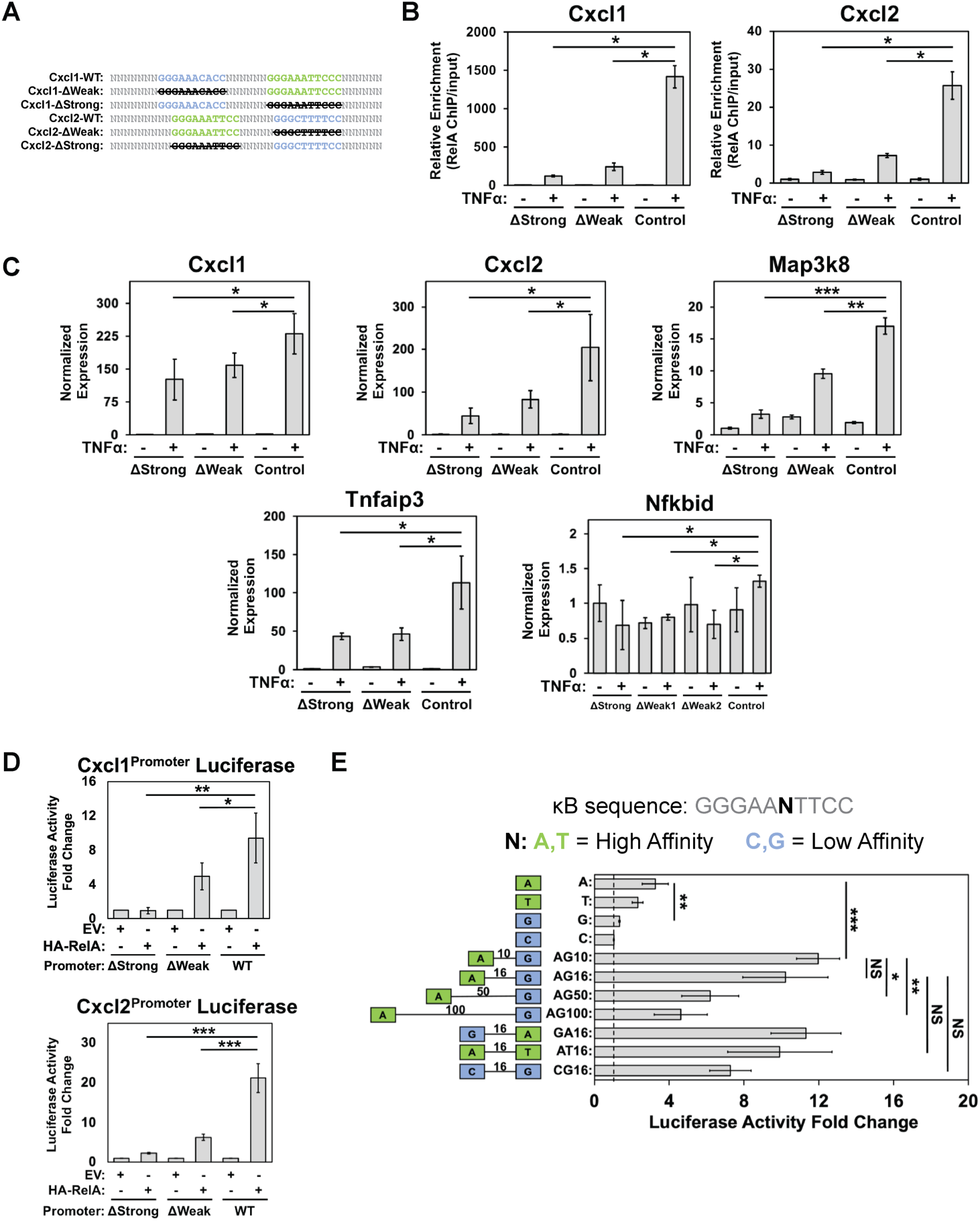
Multiplicity of κB sites (strong and weak) influence recruitment of RelA and transcription. **A.** A schematic of the arrangements of strong and weak κB sites in Cxcl1 and Cxcl2 promoter regions targeted individually by CRISPR-mediated deletion. Separately, a nearby off-target site of the promoter was also mutated using CRISPR to be used as a gene-specific control. **B.** Effect of deletion of strong, weak, or a non-κB site (control) of Cxcl1 and Cxcl2 promoters on RelA ChIP-qPCR upon 30-min TNF-α stimulation. Data is normalized by amount of input DNA, and changes represented is relative to promoter with deletion of control non-κB site. *P < 0.001. P value was calculated by two-tailed Student’s t-test. n = 3 independent experimental replicates. Error bars represent standard deviation. **C.** Normalized transcript levels of Cxcl1, Cxcl2, Tnfaip3, Map3k8, and Nfkbid in MEF cells upon 1 h TNF-α stimulation measured by RT-qPCR following deletion of a strong, weak, or a control site. Data represented are normalized by Gapdh expression. *P < 0.05, ** P < 0.001, *** P < 0.0001. P value was calculated by one-tailed Student’s t-test. For all data points, n = 3 experimental replicates. Error bars represent standard deviation. **D.** Expression of luciferase reporter plasmids driven by Cxcl1 or Cxcl2 promoter (wild-type or mutant) regions in HEK 293T cells co-transfected with HA-tagged RelA or control HA-tagged EV. Luciferase readings are normalized with control Renilla luciferase activity, and fold-change is relative to EV for a given luciferase construct. *P < 0.05, **P < 0.01, *** P < 0.0001. P value was calculated by one-tailed Student’s t-test. For all data points, n = 3 experimental replicates. Error bar represent standard deviation. **E.** Expression of luciferase reporter plasmids under various synthetic promoters (containing single or combination of differentially placed strong and weak κB sites) in HEK293T cells co-transfected with HA-tagged RelA expression vector. Numbers within schematic on left represent number of bp separating κB sites. Luciferase readings are normalized to control Renilla luciferase activity, and fold change is relative to control EV for the given luciferase construct. *P < 0.05, **P < 0.01, *** P < 0.005. P value was calculated by one-tailed Student’s t-test. For all data points, n = 3 experimental replicates. Error bars represent standard deviation.

To further support the authenticity and functionality of these weak sites in transcription, we assessed expression potential of promoter segments from Cxcl1 and Cxcl2 genes (as used in EMSA in Supplementary Figure 1D) using luciferase reporter constructs. Mutation of either the strong or the weak site led to a drastic reduction in reporter expression, similar to that observed in the cellular context. This provides further support to both strong and weak κB sites of the Cxcl1 or Cxcl2 promoters being critical in optimal expression, and allude to a functional synergy between the multiple sites during transcriptional activation (**Figure 2D**). To analyze this synergy further, we assessed expression of reporters with synthetic promoters containing various combinations of strong and weak κB sites with different spacing and orientations. We not only observed that two sites act synergistically but also observed a strikingly robust reporter activity from two weak sites similar to that observed with two strong sites (**Figure 2E**). Reporter activity decreased progressively as the spacing between two sites is increased to 100 bp. Overall, these observations reveal a strong activating influence of clusters of κB sites, strong or weak, in enhancing gene expression – outperforming the expression potential of a single strong site.

### A multitude of nuclear proteins associate with κB DNA-bound NF-κB upon inducing signal

The weak affinity of most κB sites found in promoter regions of RelA-responsive genes raised queries both about their functional authenticity and the possibility of disguised processes that could provide multisite cooperativity rendering effective recruitment of RelA dimers to promoter elements *in vivo*. Since nucleosomes are known to gather a large array of nuclear factors, it is not far-fetched that additional nuclear factors could influence recruitment of RelA or other transcription factors in cells. To explore this, we investigated factors in nuclear extracts of MEF cells post 0.5-and 3-h TNF-α-stimulation that associate with Cxcl2 promoter DNA fragment (sequence mimicking that of EMSA and reporter expression assays) biotinylated for pulldown assays. The pulled-down proteins were identified by mass-spectrometry, and the specificity of factors was established by comparison of results with wild-type (WT) and a mutant (dκB; does not bind RelA since both κB sites deleted) promoter fragments (**Supplementary Figure 3A**). Several nuclear proteins (**Figure 3A**, **Supplementary Table 2**) including other transcription factors (TFs), DNA-repair proteins, RNA-binding proteins, and metabolic enzymes were observed. Various promiscuous/non-sequence specific DNA-binding proteins were also observed. Among the TFs, members of the NFAT family were particularly notable, and that of TEAD, SMAD, and CUX families were well-represented. Among the RNA-binding proteins, the SR family splicing factors and hnRNPs were abundant. Significant difference of associated factors was observed in pull-downs with nuclear extracts of MEF cells isolated at 0.5h versus 3h post-TNFα stimulation. We surmise that this post-induction temporal dependence is likely due to significantly altered characteristics of proteome. The report of a large number of RelA-interacting proteins that includes several transcription factors in the whole cell extract has recently been reported (30). To investigate if association of these nuclear factors display a dependence on promoter-characteristics, we performed similar experiments with the Ccl2 promoter DNA fragment (**Figure 3A**). The association of proteins to Cxcl2 and Ccl2 DNA fragments were largely similar, with a few exceptions. Pathway analysis revealed significant enrichment of proteins in transcriptional regulation, NF-κB signaling, and NFAT signaling pathways (**Figure 3B**). Further investigation with extracts from LPS-treated bone marrow-derived macrophage (BMDM) cells corroborated pull-down of DNA-binding TFs, RNA-binding proteins, and various enzymes (**Figure 3C** and **Supplementary Table 2**). Notably, NFAT factors were pulled down from extract of LPS-treated BMDM cells as observed with extract of TNF-α-treated MEF cells.

**Figure 3.**
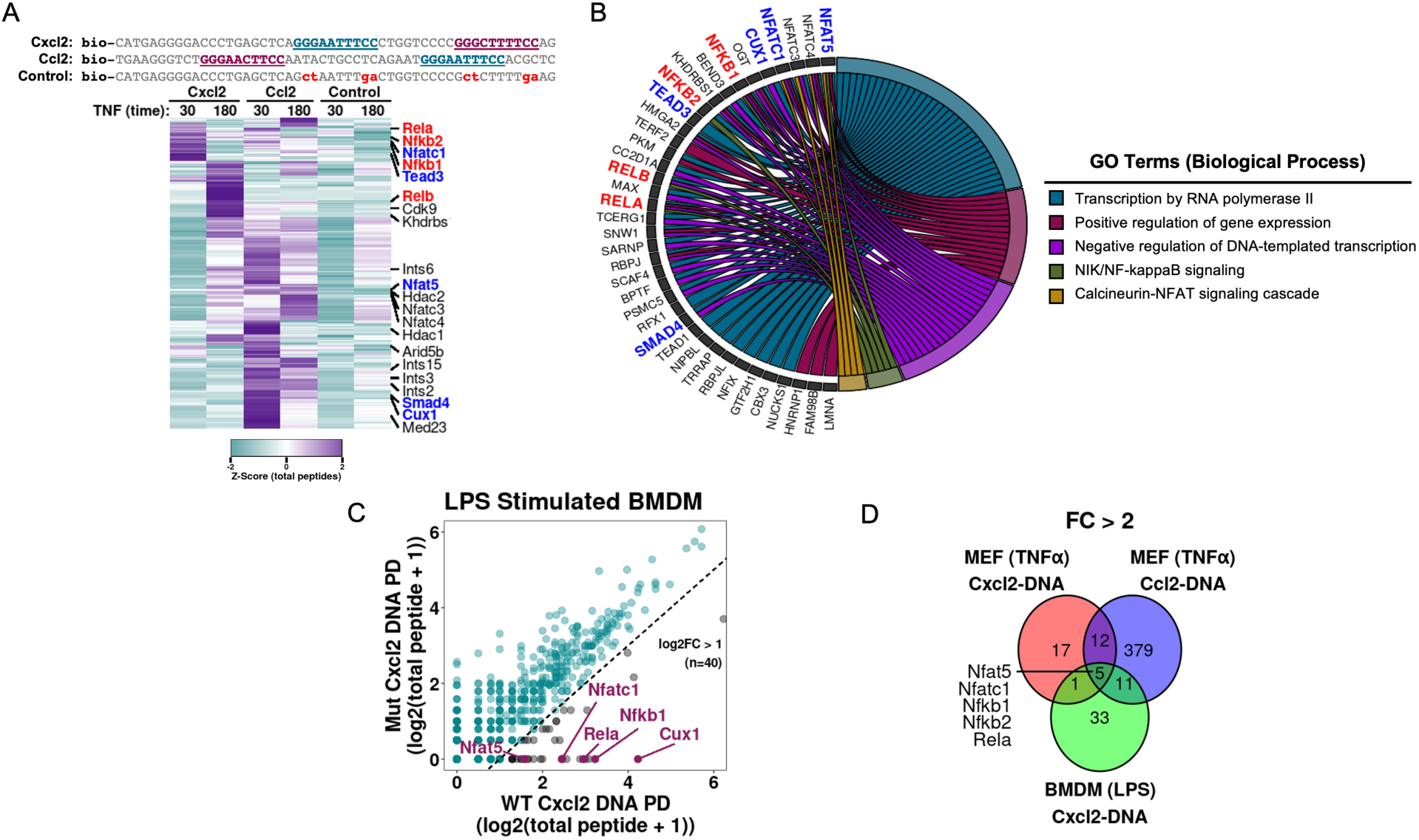
Association of a multitude of proteins to DNA adapted from tandem κB sites observed in RelA-responsive promoters of Cxcl2 and Ccl2 genes. **A.** (Top) Sequence of biotinylated DNA fragments used in pulldown experiments -containing κB sites from promoter region of Cxcl2 or Ccl2 genes, or a control DNA with no κB sites. (Bottom) Heatmap representation of total peptide (k-means clustered) Z-Scores with mass-spec of DNA-mediated pulled-down from nuclear extracts of TNF-α-treated MEF cells. Cells were stimulated with TNF-α for either 30 or 180 mins. Total peptides for each protein were normalized by total number of peptides identified in corresponding pulldown, and proteins presented represent average Z-score of Cxcl2 and Ccl2 > 0.5 relative to the control pulldown. NF-κB members are annotated in red and the putative cofactors in blue. Data represent an average of n = 2 experimental replicates. **B.** Chord diagram of biological pathway-specific gene ontology classification of enriched proteins identified in A. **C.** Correlation plot of total peptide counts from proteins identified by MS in nuclear extracts of LPS-treated BMDM enriched by binding to Cxcl2 DNA fragment compared to mutant DNA. Dotted line represents a log2FC of 1 for Cxcl2 DNA /mutant DNA. Data represent an average from n = 2 experimental replicates. **D.** A Venn diagram displaying commonality and specificity of cofactors from different cell types, DNA, and stimuli.

We explored possible roles of other transcription factors by focusing on Nfatc1, Nfat5, Tead3, Cux1, and Smad4, all of which can regulate transcriptional programs through their own cognate sequences. These factors were observed in pulldown with both Cxcl2 and Ccl2 promoter elements with cellular extracts harvested from at least one of two post-induction time points. We tested direct interaction of factors (FLAG-fused) with RelA (HA-tagged) by pull-down to ascertain if direct interaction of factors with RelA could regulate recruitment of RelA to specific promoters (**Supplementary Figure 3B**). We observed association of RelA with all five factors, albeit of different strengths. Nfatc1 interacted with RelA the strongest and it could also pulldown endogenous RelA efficiently (**Supplementary Figure 3C**). We also observed that Nfatc1 localizes to the nucleus along with RelA upon TNF-α stimulation (**Supplementary Figure 3D**). Using EMSA, we observed that affinity purified Nfatc1 and Tead3 from HEK293 cell extract can bind to the wild-type Cxcl2 DNA probe (2WT), which contains both strong and weak κB binding sites. Binding was abolished when both sites were mutated (2DM) — similarly to that observed with NF-κB p50:RelA heterodimer (**Supplementary Figure 3E**). We further performed EMSA to assess if p50:RelA heterodimer can form a ternary complex with Nfatc1 or Tead3 on DNA. Observation of smeary, shifted bands together with a reduction of individual Nfatc1 and p50:RelA single complexes (**Supplementary Figure 3F, left**) suggests a possible formation of new complexes. Ternary complexes were not observed with fragments containing only one κB site, either strong or weak, although Nfatc1 could still bind to both of these mutant probes singly. In contrast, no new bands were detected when Tead3 was included with p50:RelA on any of the three DNA probes. Surprisingly, Tead3 bound to the WT and mutant probe containing only the strong site, but not to the probe containing only the weak site - suggesting an overlap of Tead3 consensus site with only the strong κB site (**Supplementary Figure 3F, right**). These findings suggest the possibility of Nfatc1 assembling on the Cxcl2 promoter on its own or simultaneously with NF-κB, in particular when multiple κB sites are present. In contrast, Tead3 appears to engage with the strong κB site, and the access appears to be competed out by NF-κB. Whether observed simultaneous interactions are cooperative or competitive or both and if these are a general feature across other NF-κB target promoters remain to be probed. We speculate that a combined effect of cooperative and/or competitive interactions among RelA and RelA-promoter associated TFs (termed TF-CoFs) facilitate recruitment of RelA to promoters.

### TF-cofactors support recruitment of NF-κB RelA to promoter

To test if binding of RelA to the κB sites is influenced by Nfatc1, Nfat5, Tead3, Cux1 and Smad4, we created shRNA mediated MEF knockdown (KD) cell lines. A general reduction of approximately 60-80% in protein levels was observed in KD cells when compared with a control MEF cell (scramble-KD) that underwent a similar KD procedure with a scrambled sequence (**Supplementary Figure 4A**). The amounts of total RelA in these KD cells were similar to that observed in scramble-KD MEF (**Supplementary Figure 4B**).

To examine the influence of these TF-CoFs in genome-wide binding pattern of RelA, we performed chromatin immunoprecipitation followed by sequencing (ChIP-seq) of TNF-α stimulated (0- and 30-min) KD MEFs with a RelA-specific antibody (GEO series GSE287359, **Supplemental Table 3**). Analysis of the duplicate (biological) datasets identified 9096 TNF-α-inducible (at 30-min) RelA peaks (p-value < 1E-5) in these samples (**Figure 4B**, left). Consistent with earlier reports, the majority of the RelA binding sites in the scramble-KD control cells were observed in the intergenic regions (40.9%) or gene bodies, with only a small percentage (∼11%) in the promoter region (defined as spanning -1000 to +100 bp region of the TSS) (**Figure 4B**, right). Reduced binding of RelA was observed at several sites in the KD MEF cell lines, although enhancement was also observed at a few sites. Overall, the analysis of changes in RelA binding at the TNF-α-induced peaks revealed greatest perturbation by knockdown of Nfat5 and Cux1, followed by that of Nfatc1 and Smad4 (**Figure 4C**, **Supplementary Figure 4C**). Depletion of Nfatc1 and Nfat5 revealed a common change in pattern of RelA recruitment suggesting their similar binding or competitive modalities. Knockdown of Tead3 affected binding of RelA to only a few promoters, and rather enhanced recruitment of RelA at various sites suggesting a possible role of Tead3 as a competitive repressor of RelA. When RelA ChIP-seq signal is normalized over all identified peaks, Tead3 depletion indicated a minimal effect and Cux1 depletion had a maximal effect with the rest in between (**Figure 4C**). Overall, recruitment of RelA binding was affected at a significant number of sites in at least one of the co-TF KDs, with recruitment at a few promoters being perturbed by all five KDs. Genome browser tracks revealed significant reduction of specific peaks at known NF-κB target genes - *Rgs19 (in* Nfatc1-KD), *Ccl8 (in* Nfat5-KD), and *Cd44* (in Cux1-KD*) and Cxcl2 (in* Nfat5-, Cux1-, and Smad4-KD) cells (**Supplementary Figure 4D**). Furthermore, we observed that binding of Cxcl2 promoter DNA is significantly lower to NF-κB from TNF-α-stimulated nuclear extracts of Nfatc1-KD cells than NF-κB from control nuclear extract, suggesting that differences in promoter binding can be captured *in vitro* (**Supplementary Figure 4D**). Further ChIP-based assessment in WT cells could provide a clearer understanding of the correlation of binding of Nfatc1 or Cux1 to promoters of genes that showed reduced RelA binding.

**Figure 4.**
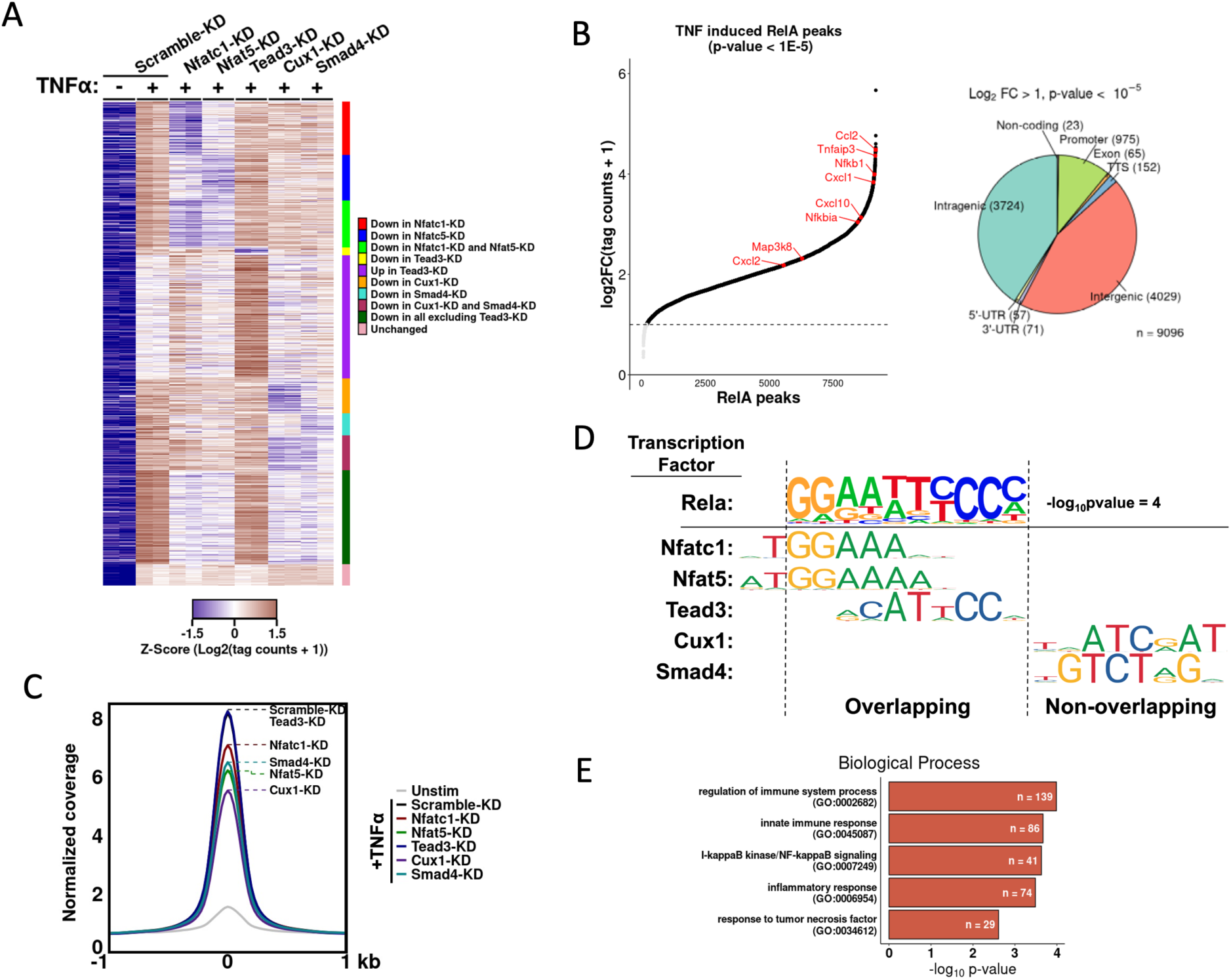
ChIP-seq indicating involvement of TF-cofactors in NF-κB RelA recruitment. **A.** Heatmap representation of genome-wide RelA binding Z-score from ChIP-Seq in TNF-α stimulated (30-min) knockdown cell lines. Peaks were filtered by TNF-α induced change in signals (log2 fold change > 1, p value < 0.05, n = 2078 peaks in Scramble-KD cells). Colors represent Z-score for each peak and categories of clustering are listed on the right. n = 2 biological replicates for all samples. **B.** (Left) Plot of ranked RelA-induced peaks in scramble-KD MEF cells following 30-min of TNF-α stimulation versus magnitude of induction relative to unstimulated cells. Dotted line represents a log2 fold change cutoff of 1. (Right) Venn diagram representing distribution of ChIP-seq peaks at various genomic loci. **C.** Profile plot showing RelA ChIP-Seq read density over all TNF-α induced peaks in knockdown cell lines. A window of +/- 1000 bp around the peak center is displayed. Peaks positions for different KD cell lines are marked in the plot using dashed lines. **D.** Motif enrichment analysis identified the κB sequence consensus as the top motif. Listed below are the consensus motifs for relevant cofactors. The Nfatc1, Nfat5, and Tead3 motifs overlap partially with the NF-κB consensus, whereas Cux1 and Smad4 motifs do not. **E.** Significantly enriched terms from gene ontology analysis of Biological Process pathways of genes with RelA-induced peaks.

We further investigated the possibility of these DNA-binding TFs influencing recruitment of RelA through binding their own DNA response elements that are either overlapping with the κB site(s) or present nearby. Motif analysis revealed that consensus κB sites were the primary TF binding sites (log p-value = -2.8E4) in control MEF cells (**Figure 4D**) and these were associated with the genes involved in the inflammatory pathways (**Figure 4E**). Tead3, Nfatc1, and Nfat5 consensus sequences were enriched at cross-linked sites; however, that of Cux1 and Smad4 were not (**Figure 4D**). Since NF-κB binding sites display significant overlap with Tead3, Nfatc1, and Nfat5, it is challenging to ascertain if these influence RelA-recruitment by binding directly to κB sites. Furthermore, Cux1 and Smad4 appear function in RelA recruitment through protein-protein interactions likely, if not exclusively by direct RelA-interaction. Overall, these results suggest that certain cofactors could support recruitment of RelA to selective promoters through their direct interaction with RelA, although TF-cofactors with overlapping sites could certainly compete and affect recruitment of RelA at other promoters.

### TF-cofactors regulate gene expression by RelA

To test if a reduced level of abovementioned TF-cofactors alter recruitment of RelA to promoters, we performed genome-wide bulk RNA sequencing of the control and five KD MEF cells (used in the ChIP experiments) post TNF-α-induction of 1h (GEO series GSE287360, **Supplementary Table 4**). Analysis of triplicate (biological) datasets revealed that TNF-α-induction increased expression of 250 genes in control scramble-KD cells (log2FoldChange > 1, p-value < 0.01), with many linked to immune response and cytokine activity pathways - as expected. Further analysis revealed genes that were differentially expressed in the TNF-α-induced KD cells when compared to TNF-α-induced control cells (p-value < 0.05; **Figure 5A**). These genes were arranged in order of highest to lowest transcript levels measured in TNF-α induced control (scramble-KD) cells, and displayed along with the transcript levels observed in the five KD cells using a heatmap (**Figure 5A**). The expression patterns differ among the KD cell lines, in contrast to the observance of a rather similar RelA ChIP signals in Nfat family KD cells. Altered expression of genes were most pronounced in Smad4-KD cells (128/250 genes, p-value < 0.05) despite the apparent minor role of Smad4 compared to the other cofactors in recruitment of RelA. Each KD affected a distinct set of genes, with some commonality among them. The differential expression of canonical NF-κB target genes is presented in **Figure 5B**. Expression of IL6 was affected in all five KD cells, whereas that of Nfkbia was not affected in any. The expression levels of most cytokine genes were impacted by KD of one or more of these TF-CoFs. These results allude to a distinct regulatory makeup of individual RelA-regulated promoters.

**Figure 5.**
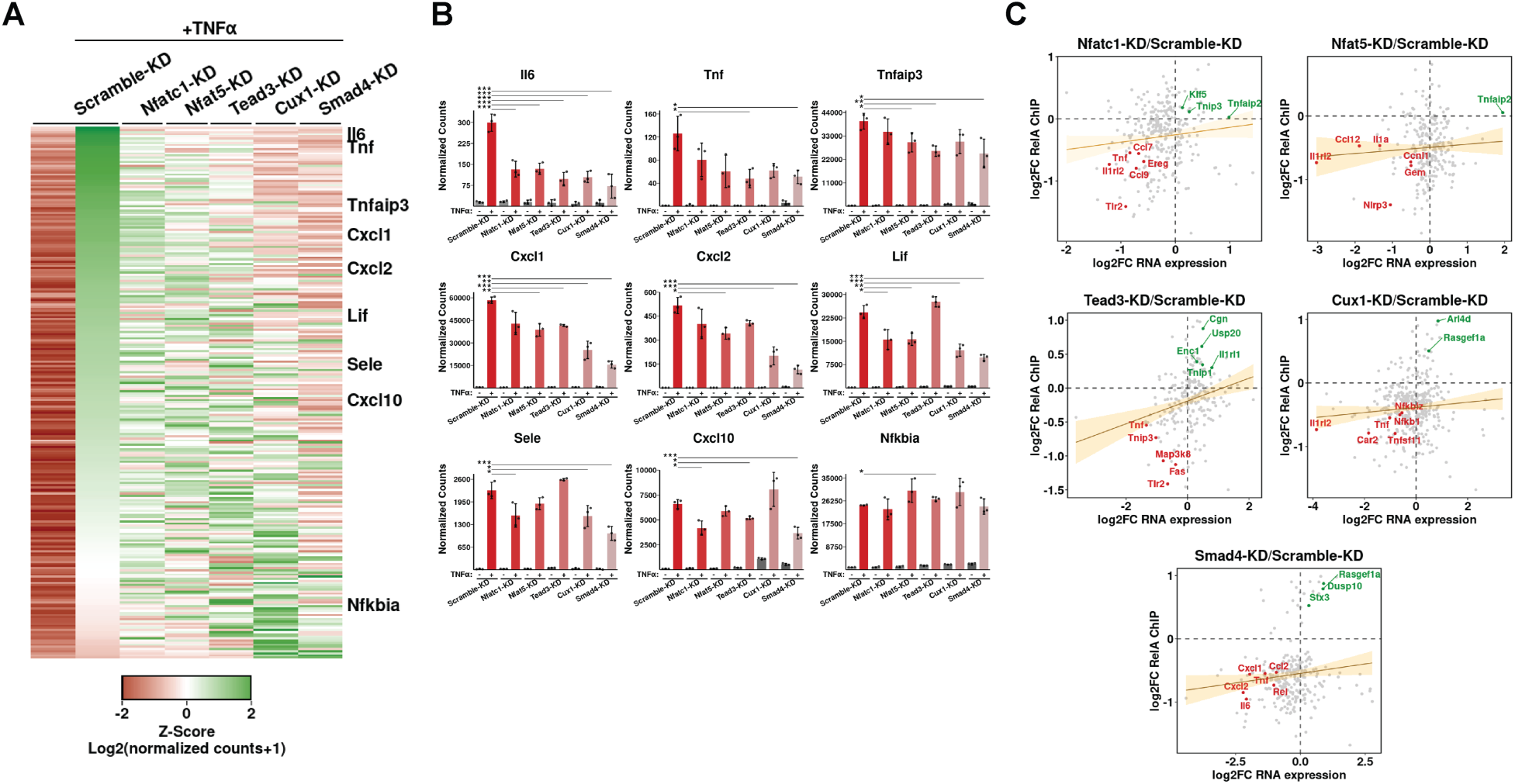
Response of TNF-α on expression of genes in various KD cell lines by RNA-seq based transcriptomic analysis. **A.** Heatmap representation of normalized Z-scores of transcript count for TNF-α induced genes in KD cell lines (log2 fold-change > 0.5, p value < 0.01, n = 250 genes in uninduced versus TNF-α treated Scramble-KD cells). Cells were stimulated with TNF-α for 1 hour before RNA isolation. Genes are sorted based on highest to lowest Z-scores of differential expressions in Scrambled-KD sample. Data represented are average of n = 3 replicates for each condition. **B.** Normalized counts of nine well-characterized RelA target genes in different KD cells presented as bar plots. Error bar represents standard deviation. n = 3 independent experimental replicates. *P < 0.05, **P < 0.01, ***P < 0.005. P value was calculated by Wald test. **C.** Plot of ChIP-seq to RNA-seq correlation in TNF-α-treated KD MEF cells compared to Scramble-KD control cells. The plots display log2 fold change of normalized transcript counts against log2 fold change in ChIP-Seq binding intensity (p value < 0.05) for TNF-α induced genes (log2 fold change > 0, p value < 0.05 in control cells) for CoF-KD relative to Scramble-KD control cells. Plots represent average of triplicate RNA-Seq and duplicate RelA ChIP-Seq data. A linear regression line is shown in orange and shaded area indicates 95% confidence interval.

The changes in transcript levels does not appear to be correlated with changes in ChIP-seq scores in KD cells (**Figure 5C** and **Supplementary Figure 5A**); however, the extent of a ChIP score reduction in KD cells did not match with the extent of transcript reduction. For example, in Nfatc1-KD cells, we observe that a higher percentage of genes with reduced ChIP-seq score also had reduced transcript levels (160/210 genes with reduced RelA binding). In this KD, ChIP-seq scores were also enhanced for several genes (83/293) of which 18 displayed a higher expression. It is possible that the observed difference in effect among genes, i.e., increased or decreased ChIP-seq scores or transcript level, may be stemming from multi-modal effects of these TF-CoFs – some cases binding to RelA and help its recruitment, and in some cases their direct engagement to overlapping or close by promoter elements, thereby antagonizing binding of RelA. A higher transcript levels of many genes with corresponding higher RelA ChIP scores were observed in Tead3 KD cells supporting a role of Tead3 in acting as a repressor of RelA (57/74 genes with increased RelA binding) (**Figure 5C**). We analyzed if the affected genes in KD cells are part of specific biological functions and observed no noticeable associations. Since no distinct correlation in changes in RelA ChIP and transcript levels among KD and control cells are observed, this could imply that RNA levels being influenced by other post-transcriptional processes (28). Overall, these results represent a rather complex relationship of TF-CoFs and recruitment of RelA.

### Clustered κB sites in the promoter/enhancer region of Cxcl1 facilitate transcription along with TF-cofactors

We explored the possible combinatorial role of clustered weak κB sites and TF-CoFs in supporting recruitment of NF-κB to promoter using the Cxcl1 gene as a model system. Analysis of 500-bp region around the transcription start site (TSS) of Cxcl1 gene (**Figure 1A**) revealed a 150-bp segment containing at least five κB sites, both strong and weak. The strongest κB site is located 57-bp upstream of TSS. RelA ChIP-seq results also revealed another peak area approximately 15 kb upstream of the TSS containing a strong (GGGATTTCCC; Z-score 8.0) and four weak κB sites within a ∼150 bp segment. Separate ChIP-seq analyses also indicate binding of RelA to this distant region and ENCODE demarcates it as a putative enhancer (**Supplementary Figure 6A**) (28).

We assessed functionality of these κB sites within the promoter and enhancer regions of the Cxcl1 gene, using luciferase reporter assay. The reporter constructs were under control of wild type (WT) or mutant (MT) promoter(P)-enhancer(E) hybrid elements (**Figure 6A**) where the promoter and enhancer are separated by a 700 bp spacer adapted from an upstream region of Cxcl1 promoter. The WT construct (E^WT^-P^WT^) exhibited ∼20-30-fold enhanced reporter activity compared to the mutant construct (E^MT^-P^MT^) with all κB site mutated (**Figure 6B**). The expression with construct containing mutated enhancer κB sites (E^MT^-P^WT^) was reduced to ∼ 30%, whereas that with construct containing mutated promoter κB sites (E^WT^-P^MT^) reduced to ∼ 10%. The expression construct driven by strong sites of both the promoter and enhancer elements (E^S^-P^S^) led to a 3.5-fold increase, i.e., ∼10% of the activity from E^WT^-P^WT^ construct whereas that driven by just the <u>s</u>trongest κB <u>s</u>ite of the promoter (E^MT^-P^S^) displayed only a 2-fold increase in luciferase activity. These results strongly suggest 1) a cooperation between the enhancer and promoter elements and 2) a potent role of weak κB sites in boosting functionality of strong sites present in both the promoter and enhancer regions (**Figure 6B**).

**Figure 6.**
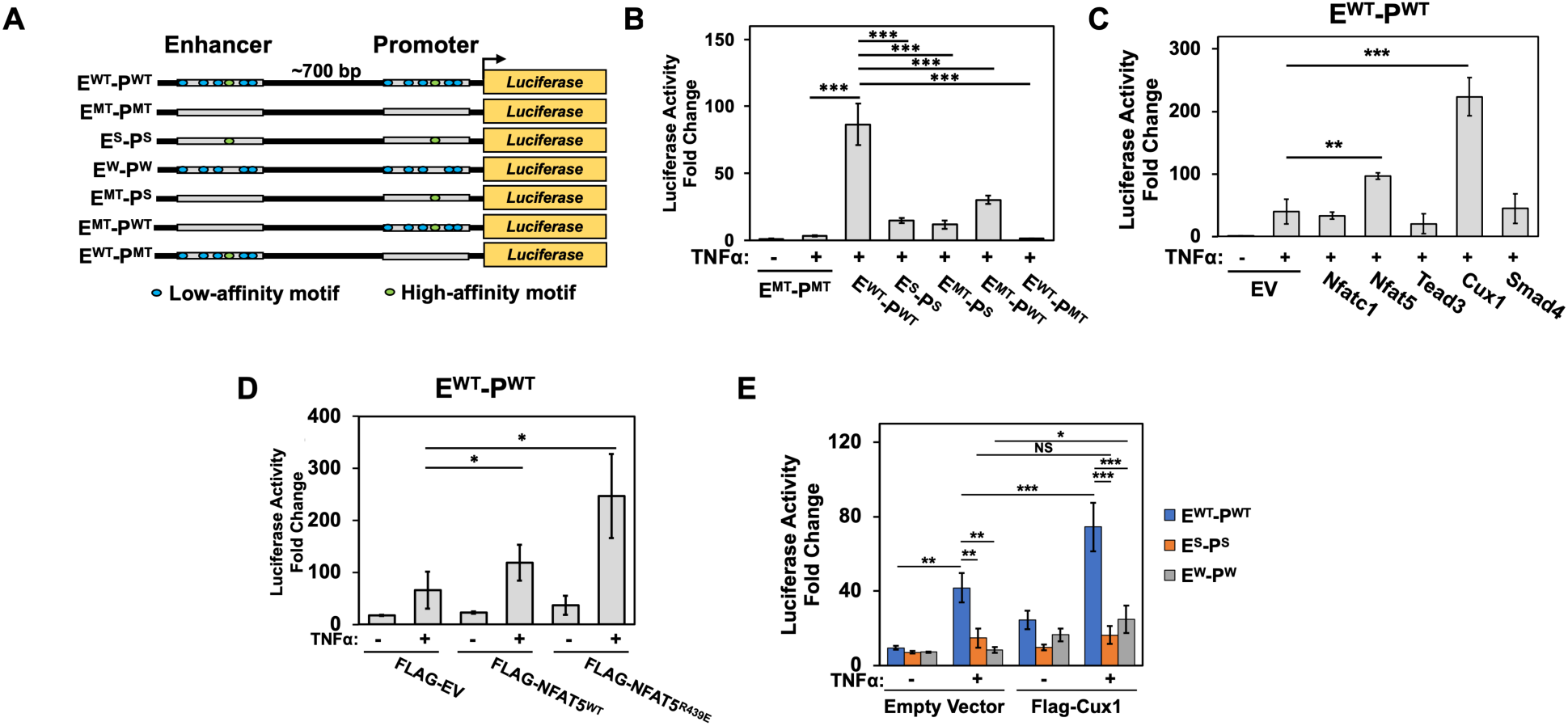
Clustered κB sites at the Cxcl1 enhancer and promoter regions regulate RelA-mediated transcriptional activation. **A.** A schematic of WT and mutant luciferase reporter constructs under Cxcl1 regulatory regions (enhancer and promoter) containing multiple κB sites. The core enhancer and promoter regions are separated by a 700 bp fragment situated upstream of the promoter and downstream of the enhancer. Green and blue circles represent high- and low-affinity κB motifs. **B.** Plot of luciferase reporter activity (see Methods) from various Cxcl1 enhancer-promoter hybrid constructs assayed in HeLa cells. **C., D.** Plot of luciferase activity from Cxcl1 E^WT^-P^WT^ luciferase construct cotransfected with control EV or Flag-tagged cofactors in HeLa cells. Fold changes are relative to unstimulated E^MT^-P^MT^ activity. **E.** Plot of Cux1-dependent luciferase activity from Cxcl1 luciferase constructs containing both strong and weak sites (blue), strong sites only (orange), and weak sites only (grey) in HeLa cells. Fold changes are relative to unstimulated E^MT^-P^MT^ activity.

Next, we assessed expression potential of these reporters under conditions where the putative TF-cofactors were overexpressed. Overexpression of Nfat5 caused a significant enhancement in reporter expression, which was much more enhanced upon overexpression of Cux1; however, overexpression of Nfatc1, Smad4, and Tead3 displayed no discernible effect (**Figure 6C**). Previously, we observed a reduction of RelA ChIP-seq scores in all TF-cofactor KD cells except for Tead3. It is possible that use of truncated Cux1 and Nfatc1 (Cux1^762-1516^ and Nfatc1^1-698^) rendered these as activator or repressor, respectively. To further assess if TF-cofactors could be functioning without requiring their DNA binding activity, we compared reporter activity from E^WT^-P^WT^ construct upon overexpression of either WT Nfat5 or its DNA-contacting residues mutated mutant (RY219SS). A slight enhancement in reporter activity was observed with mutant Nfat5 compared to WT (**Figure 6D**). We verified that the corresponding mutation in recombinant Nfatc1 (R439E) mutant does not bind DNA by EMSA, whereas neither the recombinant WT nor mutant Nfat5 RHR displayed observable binding by EMSA, perhaps due to distinct binding mechanisms (Nfatc1 binds DNA primarily as a monomer and Nfat5 as a dimer) (**Supplementary Figure 6C-6E)**). We also examined the effect of Cux1 on reporter activity from both weak (E^W^-P^W^) and strong (E^S^-P^S^) constructs. Although both mutants displayed a marked reduction in reporter expression, the presence of Cux1 significantly activated reporter expression from the E^W^-P^W^ construct containing only the weak sites. Its effect on the E^S^-P^S^ was not statistically significant (**Figure 6E**). None of these weak sites bear resemblances to Cux1 consensus sequence. These results support the possibility of a TF-cofactor, such as Cux1, likely enhancing recruitment of RelA using its non-DNA-binding activity and acting primarily through weak κB sites. Collectively, these data hints at a collaboration of κB sites and a large set of TF-cofactors in recruiting NF-κB to enhancer/promoter regions.

## DISCUSSION

In this study, we observed that promoter of genes rapidly activated by RelA dimers in response to stimuli often contain not one but multiple κB sites present in clusters. We explored the role of these κB sites of a broad spectrum of binding affinity in recruitment NF-κB RelA dimers to promoter and in transcription. Most κB sites within the cluster exhibited rather a weak affinity in vitro towards RelA dimers. In parallel, we also observed association of many accessory factors, including some other DNA-binding transcription factors, to promoter elements bound to NF-κB RelA. Analysis of these DNA-binding transcription factors indicate that they regulate recruitment of RelA to the transcription site using both their DNA-binding and non-DNA-binding activities.

### Clustering of weak and strong κB sites

We analyzed the presence of multiple κB sites within the promoter and enhancer regions of several rapid response murine NF-κB target genes – primarily cytokines (e.g., Cxcl1, Cxcl2, Tnfa) and NF-κB/IκB family members (e.g., Nfbia, Nfkid, Nfkb1, Nfkb2). The κB sites appeared to be clustered within segments of 150-200 bp, a length that corresponds to nucleosome-free DNA regulatory regions in activated genes (29). These rapid response promoters usually contained one strong site with multiple weak sites clustered around it. The in vitro affinity of individual weak κB sites to NF-κB RelA homodimer or p50:RelA heterodimer are often undetectable (i.e., Kd > ∼1500 nM). These sites often deviated from the consensus at several bases. Interestingly, X-ray structural studies revealed that the global nature of protein-DNA interaction is preserved regardless of the difference in binding affinity, and despite both the protein and DNA undergoing conformational adjustments in forming the complex.

### Significance of weak κB sites

The functional significance of weak TF-binding sites has been recognized, particularly in regulation of developmental genes of lower eukaryotes (18,21,22). However, their mechanistic role(s) in facilitating formation of transcription complexes remain unclear. We explored this significance by mutating weak sites in vicinity of the putative strong site in promoter of five rapid response RelA target genes involved in the non-developmental programs. In all cases, we observed diminished level of RelA recruitment and transcription, thereby underscoring a functional role of weak sites in the enhancer-promoter-TSS region. Reporter assays further demonstrated synergistic effects of artificially clustered κB sites in activating transcription. Since an in vitro study indicated a lack of cooperativity through tandem κB sites in recruiting RelA dimers (31), we anticipate alternative mechanisms underlying multi-site cooperativity possibly functioning not through DNA (30).

### Role of other transcription factors in recruiting RelA

We observed association of various factors, including multiple TF-cofactors, at the clustered κB site region of a native promoter and tested if they regulate recruitment of NF-κB RelA dimers. Characterization of five of these TF-cofactors indicated their interaction with RelA to be weak. However, a subset of these TF-cofactors collaborated with the κB sites in their native context to help recruit RelA at specific transcription sites. It is possible that a cofactor tags multiple RelA dimers using their different domains and facilitate recruitment of RelA to a κB cluster. Alternatively, multiple interacting cofactors can establish a protein network at a specific subset of κB sites in the enhancer/promoter of a target gene and guide RelA to that region.

Some TF-cofactors (e.g., Cux1 and Smad4) appear to function without requiring their own DNA binding since no enrichment of their binding sites were observed at RelA bound genomic regions. However, it is possible that these cofactors could regulate RelA-recruitment through contacting DNA. This is particularly true when DNA recognition sites of TF-CoFs overlap with κB sites. To test if DNA-binding of Nfat5 influences recruitment of RelA, we measured transcriptional potential of a DNA-binding defective mutant of Nfat5 in a reporter assay. The mutant showed an enhancement albeit minor of the reporter expression suggesting a possibility of these TFs to regulate recruitment without involving their DNA-binding activity. This observation agrees with reports indicating recruitment of TFs to genomic region lacking their cognate DNA sites (32). In contrary, the removal of Tead3 enhanced gene expression, consistent with its competition with RelA for strong κB sites binding. This is expected since its recognition site is closest to κB sites among the five TF-cofactors.

### Role of enzymes and RBPs in recruiting RelA

We postulate that activated nuclear RelA is trapped by various cofactors at promoter region, thus effectively enhancing concentration of RelA and enabling its recruitment to weak sites. Cofactors not belonging to TF-cofactors perhaps facilitate recruitment of RelA by different means, such as by modifying RelA. The association of kinases, methylases, acetylases, deacetylases, and glycosylases at RelA target promoters prompts us to propose that these enzymes could be modifying both protein and DNA and regulating assembly or disassembly of transcription complex at the start site (33) (34,35) (36,37). The RNA binding proteins may simultaneously engage to RNAs, promoter DNA and RelA to stabilize the transcription complex (38-41).

### κB site clusters in recruiting RelA

The requirement of κB sites within a cluster for recruiting RelA is unclear. Our findings demonstrate that cooperativity through protein-protein interactions between two NF-κB dimers bound to tandem sites is not a general mechanism for RelA recruitment. Although such cooperative mechanisms have been demonstrated for some transcription factors, particularly those that bind weakly as a monomer and require a supporting factor for stable binding, this does not appear to be the case for RelA (42). Although we categorize weak κB sites as a single class, their sequences and possibly their affinities for RelA vary substantially. For instance, deletion of a single weak κB site among three in the Nfkbid promoter led to only a partial reduction in both RelA binding and transcriptional activity. If all κB sites were equally important and acted simultaneously and cooperatively, we would have observed more pronounced defects. As shown in figure 1A, the promoter and possible enhancers of rapidly and strongly activating NF-κB genes that have more than three weak κB sites. At least in case of the Cxcl1 promoter-enhancer, the collective activity of the ∼10 weak κB sites (the precise number of sites that can engage RelA is not experimentally assessed) is comparable to that of two strong κB sites. Transcriptional activation from the weak κB sites appears to require Cux1, unlike that from the strong sites, supporting a role of Cux1 in recruitment of RelA to promoter with weak sites. Importantly, cofactor engagement appears to be stimulus specific (different cofactors are activated by TNFa and LPS), and temporally regulated, suggesting a model of dynamic cooperativity for transcriptional synergy. This dynamic and flexible regulatory mechanism offers a broader range of control than a rigid cooperative model. It also implies that single bp changes can modulate activity, contributing creating a malleable complex finely tuned by both protein and DNA modifications (27,43).

### Multi-site and multi-factor-based recruitment of a transcription factor to enhancer-coupled promoter

We observed a strong contribution of weak TF-binding sites present in the enhancer and the promoter regions through reporter expression assays. This revealed intriguing possibilities of a multisite collaboration in facile recruitment of a TF. We speculate that clustered sites within both the enhancer and promoter regions capture multiple copies of the target TF in association with multiple other TFs. Central to this model is the idea that the TF triggering activation is stabilized at the promoter with help of a dynamic condensate that include juxtaposed promoter and enhancer regions enabled through chromatin looping and a large body of multiple factors. This enhancer–promoter communication provides a much greater combinatorial possibilities of protein–protein interactions for a finer regulatory control compared to a TF binding to tandem sites in a linear promoter region. This is particularly significant in context of promoters and enhancers enriched with weak TF binding sites, which cannot stably recruit transcriptional machinery on their own but provide a critical support for TF binding to strong sites. A cartoon highlighting proposed formation of essential bridges through weak interactions among multiple sites and multiple factors supporting DNA-bound TF RelA for a robust and timely transcriptional output is presented in **Figure 7**. We surmise that this principle is likely part of other stimulus-induced transcription programs, where dynamic enhancer-promoter interactions, cofactor activities, and coordinated TF binding together shape the transcriptional response with high specificity and flexibility. It should be noted that alternative mechanisms of transcription factor (TF) recruitment are also possible where the promoter containing one or more strong binding sites of a single TF or multiple TFs, allowing efficient transcription with minimal to no need for enhancer or cofactors. This may be particularly relevant in sustained high-level expression of housekeeping genes.

**Figure 7.**
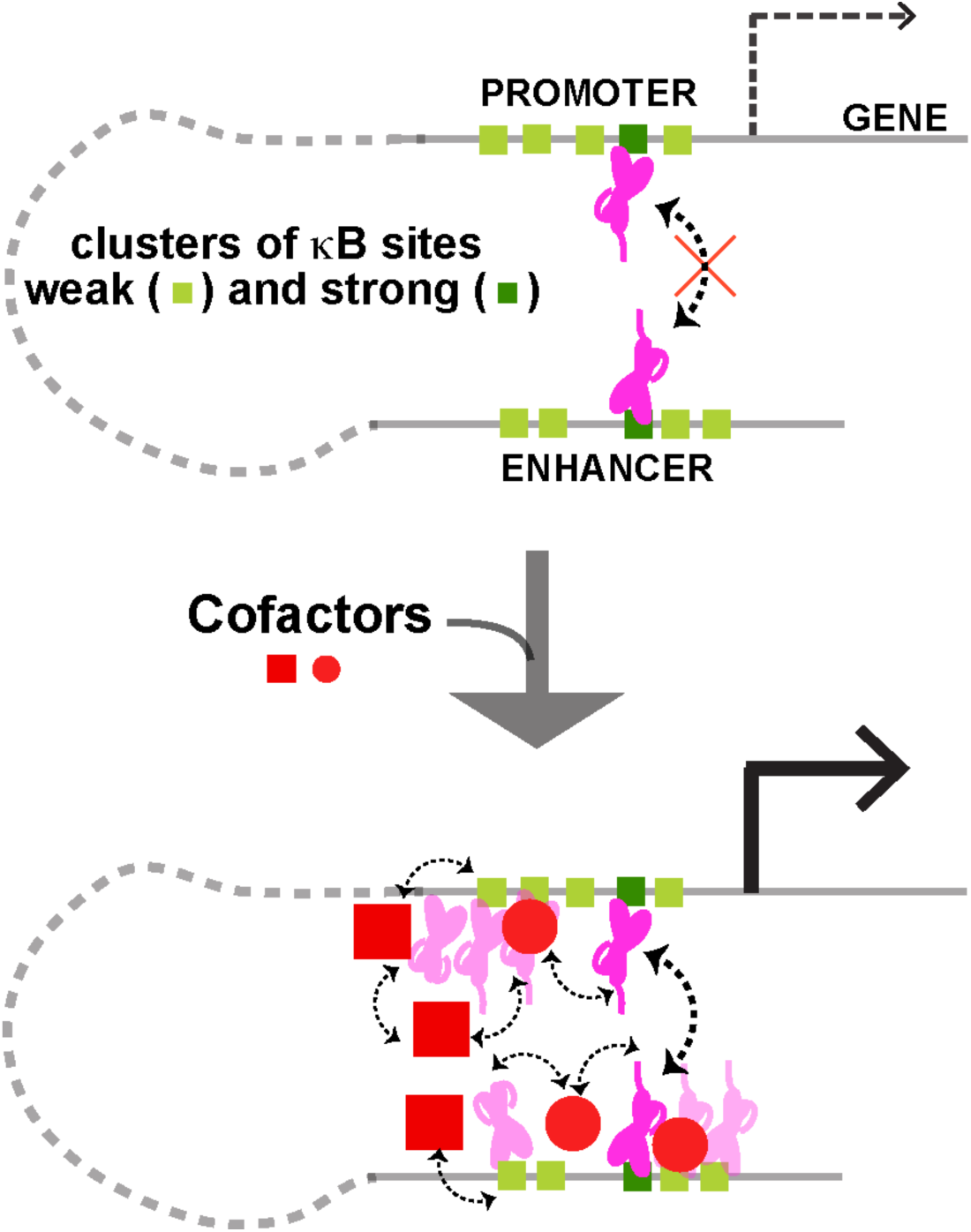
A schematic of RelA transcription complex at the Cxcl1 regulatory (promoter and enhancer) region indicating a complex interplay among strong and weak κB sites along with several cofactors. We hypothesize that interactions among multiple cofactors and NF-κB dimers within a dynamic condensate is promoted by clustered low- and high-affinity κB motifs - which eventually facilitate recruitment of NF-κB to promoter and consequently transcriptional activation. Light green and dark green squares represent low- and high-affinity κB motifs, respectively. Red circles and squares represent different cofactors. NF-κB dimers engaged to high- and low-affinity κB sites are represented in dark and light pink colors.

Overall, this study points to a significant complexity of RelA targeted promoter elements that cannot be explained by NF-κB RelA binding directly to a high-affinity cognate κB DNA. We observe cooperativity among multiple NF-κB dimers in vitro with κB sites that are spaced apart and not tandem. Our findings also allude to engagement of not only NF-κB RelA but multiple other factors including other DNA-binding cofactors – hinting at a multi-protein transcription assembly involving multiple κB sites that likely allows capture of NF-κB at the promoter even at a low concentration and activate transcription through mediator complexes and RNA polymerase II.

## Materials and Methods

The details of Materials and Methods are provided in the Supplementary Information file.

## Data Availability

Data used in this study are made available through Supplementary data file and public repositories. Mass-spectrometry data can be accessed at PRoteomics IDEntifications Database (PRIDE) under the accession PXD065004. ChIP-seq and RNA-seq data can be accessed at https://www.ncbi.nlm.nih.gov/geo/query/acc.cgi?acc=GSE287359 and at https://www.ncbi.nlm.nih.gov/geo/query/acc.cgi?acc=GSE287360, respectively. X-ray data can be accessed at https://www.rcsb.org/structure/9E6W.

## ACKNOWLEDGEMENTS

The work was supported by NIH grant GM085490 to G.G. S.S. was supported by the CMG training grant during his Ph.D., and partially supported by Siraj Therapeutics. The authors also thank the UCSD IGM Center for support with sequencing. Authors thank Alex Hoffmann of UCLA and Tom Huxford of San Diego State University for helpful discussions.

## AUTHOR CONTRIBUTIONS

S.S., T.B., and G.G. conceived original ideas. S.S. conducted most experiments. T.B, Y.S., R.S. and Y.Z. performed experiments. S.S., T.B, and G.G. analyzed data. S.S., T.B., and G.G. wrote the manuscript.

## Supplementary Information

### Materials and Methods

#### Antibodies and Reagents

The RelA (8242), H3 (9715), Lamin B1 (D4Q4Z), and Gapdh (14C10) antibodies for Western Blot and EMSA supershift assays were purchased from Cell Signaling Technology. The Nfat5 (PA1-023), Cux1 (PA5-36355), and Smad4 (MA5-15682) antibodies were purchased from Thermo Fisher Scientific. The Tead3 (A7454) antibody was purchased from ABclonal. The IκBα (0040) and tubulin (0119) antibodies were purchased from Bio Bharati Life Science. The FLAG (F1804) antibody was purchased from Sigma-Aldrich. The Nfatc1 (7294) antibody was purchased from Santa Cruz Biotechnology. The RelA antibody (C15310256) used for ChIP-Seq was purchased from Diagenode. Mouse TNF-α was obtained from Bio Bharati Life Science and used at a final concentration of 10 ng/mL for the indicated timepoints for stimulation.

#### Mammalian Cell Culture and Generation of Stable Cell Lines

WT and genetically modified mouse embryonic fibroblast (MEF), HeLa S3, and HEK 293T cell lines were cultured in Dulbecco’s modified Eagle’s medium (DMEM; Corning) supplemented with 10% fetal bovine serum (Corning), 1% penicillin streptomycin glutamine (Gibco), and appropriate antibiotics.

For generation of stable CRISPR-modified MEF cell lines, gRNA targeting a 20 nt genomic region adjacent to a NGG (N = any nucleotide) Cas9 protospacer adjacent motif (PAM) was first annealed and cloned into BsmbI (NEB) digested pLentiCRISPRv2 vector. Sanger sequencing was performed to verify cloning. The different pLentiCRISPRv2 gRNA constructs were then cotransfected with pMDLg/pRRE, pCMV-VSV-G, and pRSV-Rev expression constructs into HEK293T cells using TransIT-Lenti transfection reagent (Mirus) for generation of lentiviral particles. After 24 h, supernatant containing virus was collected and filtered through a 0.4 μm filter, and this was used to infect MEF cells at a dilution of 1:10 in the presence of 10 ng/μL polybrene (MilliporeSigma) for 48 h. MEF cells were then selected by growing in media containing 5 μg/mL puromycin. Stable shRNA knockdown MEF cells were generated similarly, except for the annealed oligos for targeting shRNA were cloned within AgeI and EcoRI sites of pLKO.1 TRC vector, which were then cotransfected with pMDLg/pRRE, pCMV-VSV-G, and pRSV-Rev expression constructs into HEK293T cells for generation of viral particles.

#### Protein Expression and Purification

Full length His-RelA (1-551) was expressed in *Spodoptera frugiperda* 9 (Sf9) insect cells using the Bac-to-Bac baculoviral expression system as previously described (12). Briefly, baculovirus infected Sf9 cells were sonicated in lysis buffer containing 25 mM Tris-HCl pH 7.5, 500 mM NaCl, 5% glycerol, 0.1% NP-40, 20 mM imidazole, 5 mM β-mercaptoethanol, and 0.5 mM PMSF. The lysate was clarified by centrifugation at 40,000g for 30 mins and incubated with Ni-NTA agarose beads (Thermo Fisher Scientific). Beads were extensively washed with lysis buffer without PMSF, and RelA was eluted with lysis buffer containing 250mM imidazole and no PMSF. Appropriate elution fractions were pooled, concentrated, and applied to a HiLoad 16/60 Superdex 200 (Cytiva) size-exclusion column equilibrated with buffer containing 25 mM Tris-HCl pH 7.5, 200 mM NaCl, 5% glycerol, and 1 mM DTT. Peak fractions were pooled, concentrated, and aliquots were flash frozen in liquid nitrogen.

For preparation of recombinant p50:RelA heterodimer, His-tagged p50 (1-435) was transformed into *E. coli* Rosetta (DE3) cells. 3-5 colonies were used to inoculate a 5 mL LB starter culture overnight at 37 °C, which was then used to inoculate 2L of LB the next day. Cells were grown at 37 °C to an attenuance (600nm) of 0.2 and induced with 0.25 mM isopropyl β-D-thiogalactpyranoside (IPTG) for 18 h at 25 °C. Cells were harvested by centrifugation at 3,000g for 10 minutes, and sonicated in lysis buffer containing 25 mM Tris-HCl pH 7.5, 250 mM NaCl, 10% glycerol, 0.1% NP-40, 10 mM imidazole, 0.25 mM PMSF, and 5 mM β-mercaptoethanol. Lysate was clarified by centrifugation at 40,000g for 30 mins and incubated in batch with Ni-NTA agarose beads (ThermoFisher), and beads were washed extensively with lysis buffer containing 20 mM imidazole and without PMSF prior to elution of recombinant His-tagged p50 in lysis buffer containing 250 mM imidazole without PMSF. The eluted p50 was concentrated and mixed with His-tagged RelA (1-551) in a 1:1 molar ratio at a final concentration of 0.1 mg/mL in buffer containing 25 mM Tris-HCl pH 7.5, 250 mM NaCl, 10% glycerol, and 1 mM DTT, and incubated for 1 h at room temperature. The p50:RelA heterodimer formed was then concentrated and separated on a HiLoad 16/60 Superdex 200 size-exclusion column equilibrated in buffer containing 25 mM Tris-HCl pH 7.5, 200 mM NaCl, 5% glycerol, and 1 mM DTT. Peak fractions of pure heterodimer were collected, concentrated, and aliquots were flash frozen in liquid nitrogen.

#### Crystallization of RelA:κB complexes and structure determination

For crystallization of RelA:κB-DNA complexes, κB-DNA duplexes were generated as previously described (1). Complexes were formed by incubating untagged RelA(19-304) at a final concentration of ∼150 micromolar with κB-DNA duplex in 1:1.2 molar ratio in buffer containing 25 mM Tris-HCl pH 7.5, 50mM NaCl, and 1mM DTT for 15 minutes at room temperature. Crystals were obtained by hanging drop vapor diffusion method at 18 °C - mixing 1μl of complex with 1μl of reservoir solution containing MES pH 5.5, 2 mM calcium chloride, 50 mM ammonium chloride, 14 % PEG3350, 1 mM spermine, and 0.05 % octyl β-D-glucopyranoside. Crystals were nucleated around the fourth day and reached a maximum size of around 50X50X200 μm around the 10th day. Crystals were soaked for 5 minutes in the mother liquor without DTT containing 21 % ethylene glycol for cryo-protection, and flash frozen in liquid nitrogen for subsequent diffraction data collection at the synchrotron source. Structure was determined by molecular replacement technique using CCP4i (2).

#### Analysis of RelA binding sites in promoter from ChIP-seq data

We analyzed previously reported RelA ChIP-Seq data sets of TNF-α stimulated MEF cells (23) to correlate genomic RelA DNA binding strength (ChIP-Seq peak intensity) with calculated motif strength (RelA motif enrichment) from established RelA-dependent target genes. The JASPAR transcription factor binding database was used for initial RelA motif identification within RelA ChIP-Seq peaks (PMC10767809). The strength of RelA binding was calculated by quantifying peak intensity at strongest RelA motif within ChIP-Seq peaks, and the strongest motif associated with a gene (defined by proximity to TSS) was considered to sort the genes. The genomic regions centered around ChIP-Seq peaks (+/- 500 bp) were analyzed for the presence of other RelA motifs of various affinities. The affinities of RelA sequence motifs defined using Z-scores were previously estimated in a SELEX study with p50:RelA (12). A Z-score boundary of 6 (equivalent to approximate in vitro binding affinity of greater than 300 nM) was used to separate strong versus weak RelA-binding κB sites. Cumulative Z-score was calculated with a dynamic window ranging from 10 to 500 bp surrounding the strongest RelA motif for an individual RelA ChIP-seq peak, and a Pearson correlation analysis between cumulative Z-score and length of analysis window was performed.

#### RNA-Seq Analyses

RNA-seq experiments were performed in triplicates. In brief, five million control or knockdown MEF cells were plated in a single well of 6-well plates (GenClone). After overnight incubation, the cells were stimulated by adding TNF-α (BioBharati) to medium at a concentration of 10 ng/mL for 1h. Cells were washed with PBS on well and treated with TRIzol (Invitrogen) to isolate RNA following the manufacturers recommendation. Isolated RNA was quantified using ND-1000 spectrophotometer (Nanodrop), its integrity was assessed with Tapestation (Agilent), and DNA libraries were prepared using 1 μg of total RNA using KAPA mRNA Hyperprep Kit (Roche) with KAPA Unique Dual-Indexed adapters (Roche). DNA libraries were PCR amplified (8 cycles) prior to analysis by Tapestation (Agilent). Next, libraries were quantified by dsDNA Qubit (Thermo Fisher Scientific), pooled, and put through 100 bases of paired-end sequencing using the NovaSeq 6000 S4 sequencer (Illumina). Sequencing quality was assessed by FastQC and high-quality reads of libraries were trimmed using Trimmomatic (v0.39) prior to being mapped onto the mm10 mouse reference genome using STAR (v2.7.10b). Transcripts were quantified using StringTie (v1.3.6) and counts were imported to R using prep.de.py3 for normalization and differential expression analysis with DESeq2 (v1.42.0).

#### ChIP-Seq Analyses

Chromatin immunoprecipitation (ChIP) was performed as previously described in biological duplicates (23). Briefly, control and knockdown MEF cell lines grown as for RNA-seq analysis were stimulated with TNF-α at a concentration of 10 ng/mL for 30 min, washed with PBS once, crosslinked by incubating with disuccinimidyl glutarate (DSG) (ProteoChem) at a concentration of 2 mM in PBS for 30 min at room temperature with gentle rocking, and further crosslinked with 1% (vol/vol) formaldehyde in PBS for 10 min at room temperature. Crosslinking reactions were quenched by incubating with 125 mM final concentration of glycine at room temperature for 5 min. Cells were washed twice with ice-cold PBS and harvested by scraping. Pelleted cells were gently resuspended in 1 mL of PBS containing 0.5% NP-40 and 0.5 mM PMSF, and incubated in a rotary shaker for 10 min at 4 °C. Nuclei were pelleted by centrifugation at 400 g for 5 min at 4 °C, washed once by resuspending in PBS containing 0.5 % NP-40, then washed with PBS, followed by resuspension in 1 mL of RIPA buffer (25 mM Tris-HCl pH 7.5, 150 mM NaCl, 0.1 % SDS, 0.1 % sodium deoxycholate, 1 % NP-40, 1 mM EDTA, 1 mM DTT, and 0.5 mM PMSF). Chromatin was sonicated using a Model 120 sonic dismembrator (Fisher) with 2 mm diameter tip with settings: 15 total cycles, 20 second pulse intervals with 40 seconds dwell time, and 80% amplitude to obtain fragments in general range of 200-1000 bp. Chromatin samples were centrifuged at 13,000 g for 10 min at 4 °C to remove debris and larger fragments. Supernatant containing fragmented chromatin of appropriate size range was diluted 1:1 with buffer containing 25 mM Tris HCl pH 7.5, 150 mM NaCl, 1 mM EDTA, 1 mM DTT, and 0.5 mM PMSF prior to immunoprecipitation steps. For immunoprecipitation, 30 μL of Protein G Magnetic beads (NEB) was mixed with 1 μL of anti-RelA antibody (Diagenode) in PBS containing 0.5 % (m/v) BSA. Beads were washed thrice with PBS before resuspending in binding buffer containing 25 mM Tris-HCl pH 7.5, 150 mM NaCl, 0.05 % SDS, 0.05 % sodium deoxycholate, 0.5 % NP-40, 1 mM EDTA, and 1 mM DTT. Sonicated chromatin was then added and samples were incubated overnight in a rotary shaker at 4 °C. Beads were washed in following buffers, with each step 2 min long at room temperature in a rotary shaker unless otherwise specified: 3X with binding buffer, 3X with buffer containing 10 mM Tris-HCl pH 7.5, 250 mM LiCl, 1% NP-40, 0.7% sodium deoxycholate, and 1 mM EDTA, 2X with buffer containing 10 mM Tris-HCl pH 8.0, 1 mM EDTA, and 0.2% Tween 20, and 2X with buffer containing 10 mM Tris-HCl pH 8.0, 0.1 mM EDTA, and 0.05% Tween 20. Beads were then resuspended in 25 μL of buffer contain 10 mM Tris-HCl pH 8.0 and 0.05% Tween 20. Libraries were prepared on beads using NEBNext Ultra II DNA Library Prep kit (NEB) following manufacturers recommendations with 0.6 μM of KAPA Unique Dual-Indexed adapters (Roche). To each reaction of 46.5 μL, we added 16 μL of nuclease free water, 4 μL of 10 % SDS, 4.5 μL of 5 M NaCl, 3 μL of 0.5 M EDTA, 4 μL of 0.2 M EGTA, 1 μL of 20 mg/mL Proteinase K (NEB), and 1 μL of 10 mg/mL RNase A (Qiagen). The samples were incubated at 55 °C for 1h for Proteinase K and RNase A digestion, followed by incubation overnight at 65 °C to reverse-crosslinking. DNA was purified using 50 μL of Ampure XP beads (Beckman Coulter) and 20 % PEG8000/2.5 M NaCl at a final concentration of 12 % PEG and eluted in 13 μL 10 mM Tris-HCl pH 8.0 and 0.05 % Tween 20. The eluted DNA was PCR amplified with KAPA HiFi 2x HotStart ReadyMix (Roche) and 10x Library Amplification Ready Mix primers (Roche) for 16 cycles. Amplified DNA libraries were then purified with Ampure XP beads at a 1:1 ratio (10 % PEG final), analyzed by TapeStation (Agilent), and quantified by dsDNA Qubit (ThermoFisher). Pooled library DNAs of a size range of 200-500 bp were selected with Pippin HT (Sage Science) and 50 bp paired-end sequenced on the NovaSeq X Plus (Illumina).

ChIP-Seq analysis was performed as previously described (3). Sequencing quality was first assessed by FastQC and libraries with high quality reads were trimmed with Trimmomatic (v0.39). Trimmed FASTQ files were mapped to the mm10 mouse reference genome using Bowtie2 (v2.5.1). Peaks were then identified using HOMER v4.11.1 (findPeaks) with the parameters “-style factor -L 0 -C 0 -fdr 0.9 -minDist 200 -size 200 -o auto”. Raw tag counts were quantified with HOMER (annotatepeaks.pl) and tag counts were averaged between experimental replicates. Motif enrichment analysis was performed using HOMER (findMotifsGenome.pl) with the parameter “-size 200”. For gene ontology analysis, induced RelA peaks passing a threshold of log2FC > 1 and p-value < 0.05 were analyzed with the gseGO function from clusterProfiler (v4.8.3) with the org.Mm.eg.db (v3.17.0). Results were annotated along the ontology of biological processes with the following parameters: nperm = 10000, pvalueCutoff = 0.05, and pAdjustMethod = ‘none’.

#### Luciferase reporter assay

Complementary primers of specific promoters were mixed at a final concentration of 2 μM in buffer containing 10 mM Tris-HCl pH 7.5, 50 mM NaCl, and 1 mM EDTA and annealed by placing them in beaker with 95 °C water that was gradually cooled to room temperature. Annealed promoters were cloned within SalI and BamHI sites of the CMXTK-Luciferase vector (obtained from Dr. D. Chakravarti, Northwestern University Feinberg School of Medicine).

HEK 293T were plated in 24-well plates 24 h prior to transfection so that they reached a confluence of ∼ 70-80 % confluence at the time of transfection. Cells were transiently transfected using PEI with indicated reporters (e.g., in Figure 2E) along with HA-RelA (1-551) or a control empty HA-vector, and Renilla luciferase expression vector under control of a CMV-promoter. Cells were harvested 24 h post-transfection by scraping in 1x passive lysis buffer (Promega), and lysate was clarified by centrifugation at 13000g for 10 min at 4 °C. Luciferase activity was measured using the Dual-Luciferase Reporter Assay System (Promega). Readings were first normalized by comparing with activity of Renilla luciferase. The effect of RelA was then calculated by comparing samples transfected with HA-RelA vs empty HA-vector. For each case, three or more biological replicates were performed, and the results are represented as mean ± standard deviations (SD).

For luciferase reporter activity using HeLa cells (e.g., in Figure 6B-E), the cells were also plated in 24-well plates 24 h prior to transfection so that they reached a confluence of ∼ 70-80 % confluence at the time of transfection. Cells were transfected using PEI with indicated luciferase reporter construct (or an E^MT^-P^MT^ construct-activity from which was later used for normalization) along with either FLAG-tagged cofactor or empty FLAG-vector (as a control) for 48 h, followed by treatment with 10 ng/mL TNF-α for 6 hours prior to harvest by scraping in 1x passive lysis buffer (Promega). Cell lysate was clarified by centrifugation at 13000g for 10 min at 4 °C. Luciferase readings were first normalized by lysate protein concentration measured by Bradford reagent. Fold-changes in activity from the luciferase reporter (compared to activity from E^MT^-P^MT^) in presence or absence of co-factor was then analyzed. Data represented are mean ± standard deviations (SD) of three or more biological replicates.

#### Electrophoretic Mobility Shift Assay

Oligonucleotides were ^32^P-end labeled with [γ-^32^P]ATP (PerkinElmer) using T4-polynucleotide kinase (New England Biolabs). Labeled DNA was purified of free [γ-^32^P]ATP using a Microspin G-25 column (Cytiva) and annealed with complementary DNA as described in luciferase reporter assay. These radiolabeled DNAs were incubated with different amount of recombinant RelA:RelA or p50:RelA at room temperature for 15 minutes in binding buffer containing 10 mM Tris-HCl pH 7.5, 50 mM NaCl, 10 % glycerol, 1 % NP-40, 1 mM EDTA, and 0.1 mg/mL sonicated salmon sperm DNA (Fisher). Recombinant proteins of different concentrations were prepared using dilution buffer containing 20 mM Tris-HCl pH 7.5, 50 mM NaCl, 10 % glycerol, 1 mM DTT, and 0.2 mg/mL bovine serum albumin for the reaction mixtures. Complexes were separated by electrophoresis at 200 V for 1 h at room temperature in a pre-ran 5% non-denaturing polyacrylamide gel in 25 mM Tris, 190 mM glycine, and 1 mM EDTA buffer. Gels were dried, exposed on a phosphor screen overnight, and shifted bands were scanned by Typhoon FLA 9000 imager (Cytiva) for analysis.

For EMSA with nuclear RelA, MEF cells were harvested by scraping in PBS and lysed in cytoplasmic lysis buffer consisting of PBS with 0.5 % NP-40 and 0.5 mM PMSF. Nuclei were pelleted by centrifugation at 300 g, washed twice with PBS, and resuspended in extraction buffer containing 25 mM Tris-HCl pH 7.5, 420 mM NaCl, 10% glycerol, 0.2 mM EDTA, 1 mM DTT, and 0.5 mM PMSF to obtain nuclear extract. Seven μg of nuclear extract (quantified by Bradford assay) was mixed with radiolabeled DNA (final concentration ∼ 2 nM) in a binding reaction of 10 μl consisting of 25 mM Tris-HCl pH 7.5, 150 mM NaCl, 10% glycerol, 1 mM EDTA, and 0.1 mg/mL sonicated salmon sperm DNA. Complexes were analyzed by non-denaturing gel electrophoreses as outlined above.

#### Real-Time qPCR of RNA samples

Total RNA of cultured cells was isolated using TRIzol reagent (Invitrogen) followed by isopropanol precipitation, and RNA concentration was determined in ND-1000 spectrophotometer (Nanodrop). cDNA was synthesized with 1 μg of total RNA using Maxima H Minus Master Mix (ThermoFisher) in a reaction volume of 5 μL. Synthesized cDNA was then diluted 4-fold with nuclease free water and 1 μL sample was used as a template for real-time qPCR using Luna Master Mix (NEB) on a Bio-Rad CFX Connect Real-Time PCR detection System. The list of primers used for RT-qPCR are listed in Supplementary Table 3. *Gapdh* was used as the housekeeping control gene for normalization. Normalized expression was calculated using the ΔCt method by subtracting the mean cycle threshold (Ct) of the target gene to the average Ct of *Gapdh*. Normalized expression was calculated as 2^-ΔCt^ from three independent experiments and are represented as mean ± standard deviation (SD).

#### DNA Pulldown Assays, Mass Spectrometry, and Analysis

For streptavidin-DNA pulldown assays, 5’ biotinylated oligonucleotides (IDT) were annealed with corresponding nonbiotinylated reverse complement oligonucleotides at a final concentration of 45 μM. For each pulldown reaction, 40 μL of 45 μM annealed DNA was mixed with 100 μL of Neutravidin agarose resin (Thermofisher) at a total volume of 400 μL in buffer containing 10 mM Tris-HCl pH 7.5, 50 mM NaCl, and 0.5 mM EDTA. Additionally, all centrifugation steps were carried out at 3,000 x g for 1 minute at 4 °C and all pulldowns were performed in biological duplicates. Biotinylated DNA was immobilized onto beads by incubation for 1 h at room temperature in a rotary shaker gently. Beads were washed extensively with pulldown buffer containing 25 mM Tris-HCl pH 7.5, 150 mM NaCl, 5 % glycerol, 0.1 % NP-40, and 1 mM DTT with a final resuspension of 50 μL per pulldown reaction. Nuclear extract of TNF-α (20 ng/mL for 30 min) treated MEF cells were prepared as previously outlined. Approximately 250 μg of nuclear extract was diluted 2.8-fold with the pulldown buffer without NaCl but supplemented with 0.5 mM PMSF to reduce the final NaCl concentration from 420 mM to 150 mM. The diluted nuclear extract was mixed with 50 μL of the biotinylated double mutant DNA (2DM) immobilized to streptavidin beads and precleared by gentle rotation at 4 °C for 2 h. Mutant beads were then pelleted and discarded, and the precleared lysate was mixed with the corresponding experimental beads overnight at 4 °C with gentle rotation. Beads were then washed 4 times with 1 mL of pulldown buffer at 4 °C. Microcapillary LC/MS/MS was performed on bead eluant for peptide identification at the Taplin Biological Mass Spectrometry Facility at Harvard Medical School using Orbitrap mass spectrometers (Thermo Fisher Scientific).

For analysis of MEF mass spectrometry data, values corresponding to total peptides identified were used. Proteins were excluded if the total peptide count was < 5 across all pulldown experiments. Values of total peptides for each protein were then normalized by dividing by sumtotal of peptides identified in each corresponding pulldown. Normalized peptides were then used to assign a Z-score value for each protein and further filtered to identify peptides that are enriched in pulldown with biotin-Cxcl2 DNA or biotin-Ccl2 DNA compared to biotin-control DNA (average Z-score of Cxcl2 and Ccl2 pulldowns > 0.5 compared to control DM pulldown). For gene set enrichment analysis, the gseGO function from clusterProfiler (v4.10.0) with the org.Mm.eg.db (v3.18.0) was used. Results were annotated along the ontology of biological processes with the following parameters: nPerm = 10000, pvalueCutoff = 0.05, minGSSize = 3, maxGSSize = 300, and pAdjustMethod = ‘none’. The chord plot was generated using the GOChord function from GOplot (v1.0.2).

#### Biolayer Interferometry Assay and Analysis

The assay was performed in Octet K2 system (ForteBio) and protocol was adapted based on Li et al (1). Briefly, biotinylated oligonucleotides were mixed with complementary oligonucleotides at a ratio of 1:1.2 in buffer containing 10 mM Tris-HCl pH 7.5, 50 mM NaCl, and 1 mM EDTA and annealed by placing samples in beaker with 95 °C water that was gradually cooled to room temperature, and then placed in ice. Hydrated Octet Streptavidin (SA) biosensors (Sartorius) were loaded with these DNA duplexes by placing them in 200 nM of DNA for 10 seconds in BLI buffer consisting of 25 mM Tris-HCl pH 7.5, 150 mM NaCl, 0.02 % Tween 20, 0.1 mg/mL sonicated salmon sperm DNA, and 1 mM DTT. A reference for background subtraction is always generated using a sensor that was not loaded with DNA. For all assays, a baseline was first obtained by incubating sensors in BLI buffer for 60 seconds. Association phase of 120 seconds was observed by dipping the sensor-biotinylated DNA probe in 12.5, 25, 50, 100, or 200 nM of recombinant His-RelA (1-551), followed by a dissociation phase of 180 seconds by placing the sensor back to BLI buffer without protein. For regeneration of DNA-bound sensor (i.e., to remove the bound protein), it was placed in BLI buffer containing 1 M NaCl for 5 seconds followed by 3X washes of 5 sec in BLI buffer. Data was processed using Sigmaplot 15 to observe the binding kinetics and obtain affinity values.

#### Sanger Sequencing

CRISPR modified MEF cell lines were harvested by scraping, and genomic DNA was collected using phenol:chloroform:isoamyl alcohol (25:24:1; ThermoFisher) and ethanol precipitation. The DNA pellet was resuspended in TE buffer and used as a template for PCR reactions using Taq polymerase (NEB) with primers targeting the Cxcl2 promoter (Supplementary Table 3). The completed PCR reaction was separated by agarose gel electrophoresis and visualized with ethidium bromide. The band corresponding to the expected 200 bp Cxcl2 promoter amplicon was excised with a sterile razor and purified using the Monarch DNA Gel Extraction kit (NEB) following manufacturer’s specifications. Approximately 10 ng of purified PCR product was then subject to Sanger sequencing (Genewiz) in both directions using either the forward or reverse primer.

#### Immunoprecipitation Assay

For FLAG pulldown assays, HEK293T cells were plated in 6-well culture dishes (GenClone) at 80 % confluence and transfected using PEI with HA-tagged RelA and either empty vector or FLAG-tagged Nfatc1, Nfat5, Tead3, Cux1, or Smad4 overexpression constructs for 24 hours. Cells were then stimulated with mouse TNF-α (BioBharati) for 30 min, washed once with ice-cold PBS, and lysed by direct application of lysis buffer containing 25 mM Tris-HCl pH 7.5, 150 mM NaCl, 5 % glycerol, 1 % NP-40, 1 mM DTT, and 0.25 mM PMSF to the well. Lysate was clarified by centrifugation at 13,000g for 15 min at 4 °C. Anti-FLAG M2 agarose beads (Sigma) were incubated with the lysate for 2 h at 4 °C in a gentle rotary shaker. Beads were washed extensively with lysis buffer, and 4x SDS gel loading dye was added to a final concentration of 1x prior to heating at 95°C in a heat block for 5 minutes. Samples were separated on a 10 % SDS-PAGE gel at 200 V for 45 mins and transferred to nitrocellulose membrane for Western blot analysis.

## Supplementary Figures

**Supplementary Figure 1.**
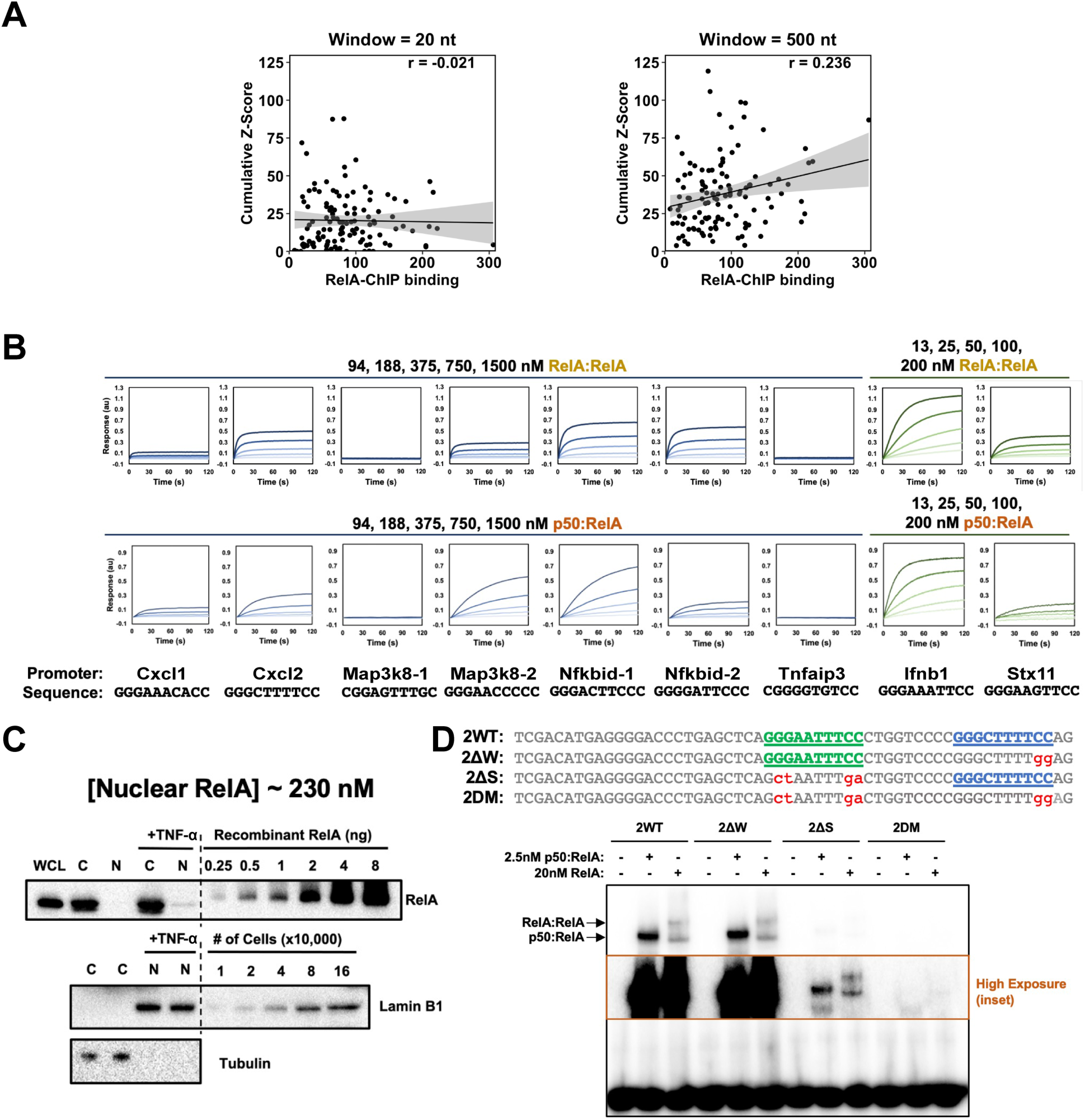
**A.** Pearson’s correlation plots between RelA ChIP-seq score and cumulative Z-score with a 20-bp window (left) or a 500-bp window (right) around the strong site. A linear regression line is shown in black and shaded area indicates 95% confidence. n = 116 peaks analyzed. **B.** The binding of κB site sequences to full-length RelA homodimer (top) and p50:RelA heterodimer (bottom) by BioLayer interferometry assay. Binding to Ifnb1 and Stx11 κB sites, present in many promoters, are used as references of affinity. Concentrations of RelA homodimer or p50:RelA heterodimer used in the assay are indicated above plots. **C.** Immunoblot analysis of RelA from whole cell lysate (WCL), cytoplasm (C), and nuclear (N) fractions of HeLa S3 cells with or without TNF-α stimulation. Purified recombinant RelA was used as a standard for quantitation of RelA amount, and Lamin B1 was used as a standard for estimating cell number. **D.** EMSA of wild-type and mutant promoter sequences derived from Cxcl2 gene with RelA homodimer and p50:RelA heterodimer. Strong and weak κB site sequences are colored in green and blue, respectively, and mutated nucleotides are denoted in red. A high exposure inset is marked with a brown border.

**Supplementary Table S1:** Crystallographic data used in obtaining the structural model of RelA:κB DNA (Cxcl2 weak) complex, and refinement statistics of the model.

|  |  |  |
| --- | --- | --- |
| **Structural Model** |  | RelA:κB DNA (CxcI2 weak T-centric) |
| **Deposition ID** |  | 9E6W |
| **Data Collection** |  |  |
| Wavelength (Å) |  | 0.97946 |
| Resolution range (Å) |  | 50.0-2.03(2.07-2.03) |
| Space group |  | P 21 21 2 |
| **Cell dimensions** |  |  |
| a, b, c (Å) |  | 118.79 133.08 45.36 |
| α, β, γ (°) |  | 90.000 90.000 90.000 |
| Total reflections |  | 604462 |
| Unique reflections |  | 46623 |
| Redundancy |  | 13.0(10.6) |
| Completeness (%) |  | 99.4(99.5) |
| I/σ(I) |  | 94.9/3.1(2.7/2.1) |
| Rpim |  | 0.022(0.547) |
| CC1/2 /(CC*) |  | 1.00(0.681)/1.00(0.90) |
| **Refinement** |  |  |
| Resolution range (Å) |  | 37.48-2.04(2.10-2.04) |
| No. of reflections (work/test set) |  | 35763/1752 |
| Rwork/Rfree |  | 0.244/0.283 |
| Number of non-hydrogen atoms |  | 5386 |
| **r.m.s. deviations** |  |  |
| Bonds (°) |  | 0.0025 |
| Angles (°) |  | 0.9609 |
| **Ramachandran plot** |  |  |
| Favored (%) / allowed (%) |  | 96/4 |
| Outliers (%) |  | 0 |
| Rotamer outliers (%) |  | 2 |
| Clashscore |  | 7 |
| Average B-factor (Å<sup>2</sup>) |  | 35 |

**Supplementary Figure 2.**
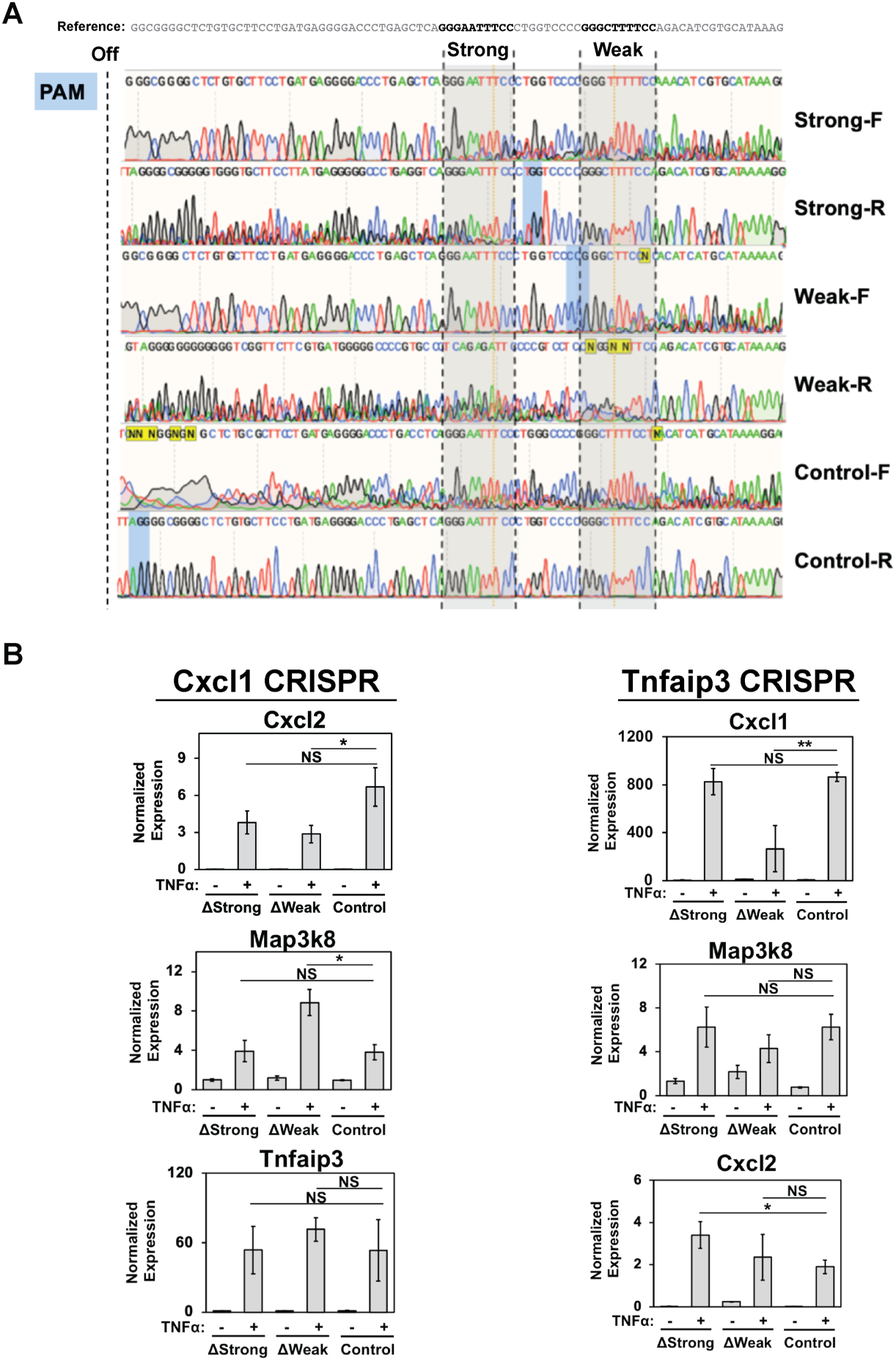
**A.** Sanger sequencing for verification of CRISPR-based mutagenesis of Cxcl2 promoter in MEF cells. Bulk cell sequencing was performed on a 200 bp amplicon of the Cxcl2 promoter region generated by PCR of gDNA from corresponding CRISPR-modified cell lines. Traces represent Sanger sequencing results in both the forward and reverse direction, with perturbations to sequencing traces indicating upstream genomic modifications. The mm10 reference sequence of the Cxcl2 promoter is shown above. The PAM sequences used to generate the three mutants are highlighted in blue shades. **B.** Normalized transcript levels of corresponding off-target genes upon CRISPR-mediated targeting of strong, weak, or control sites of Cxcl1 (left) or Tnfaip3 (right) measured by RT-qPCR. MEF cells were stimulated with TNF-α for 1 h. Data represented are normalized by Gapdh expression. *P < 0.05, ** P < 0.001, NS = non-significant. P value was calculated by one-tailed Student’s t-test. For all data points, n = 3 experimental replicates. Error bar represent standard deviation.

**Supplementary Table S2.** Total counts of identified proteins in MEF nuclear extract pulled down by biotinylated κB-DNA:RelA using mass-spectrometry. Individual results of experimental duplicates for each type of DNA used in pulldown and duration of TNF-α treatment are provided. (Table attached as a separate file denoted as SupplementaryTable2.xlsx)

**Supplementary Figure 3.**
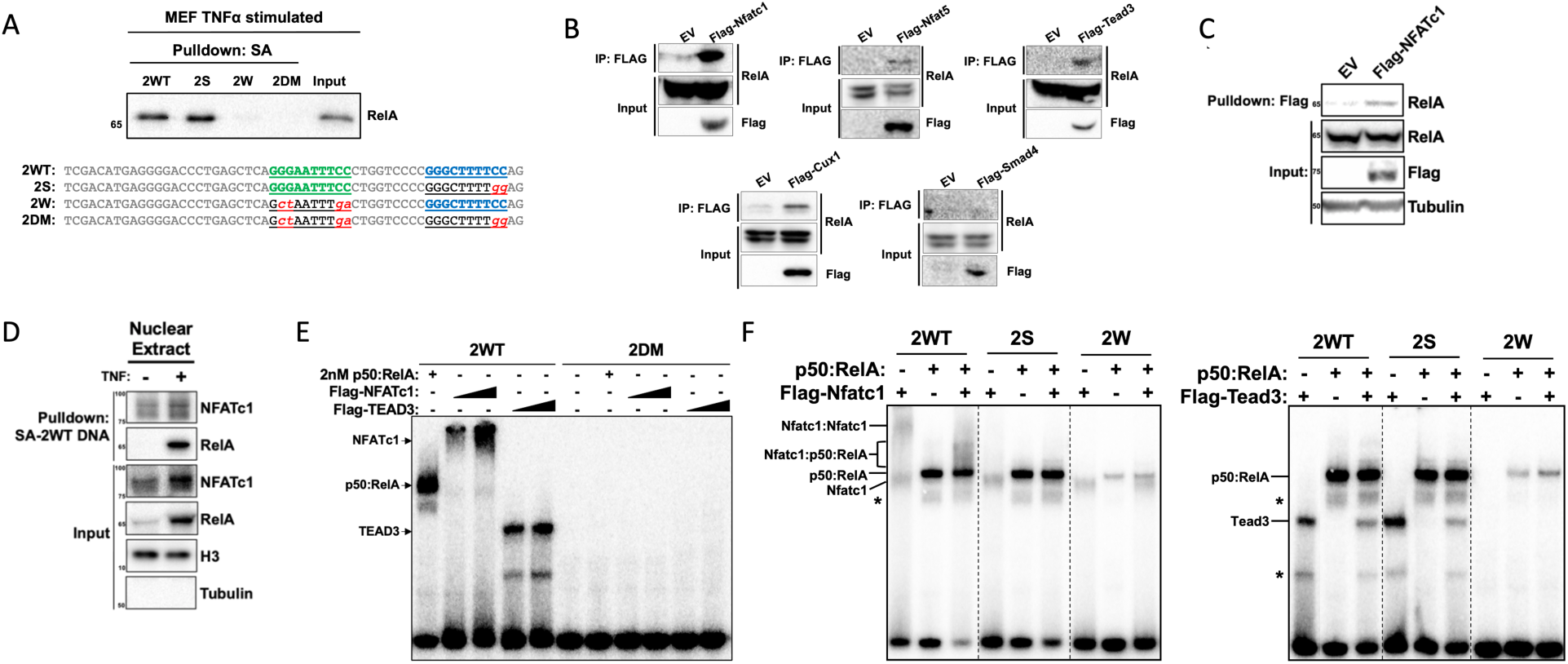
**A.** Immunoblot analysis of pulldowns with biotinylated wild-type and mutant Cxcl2 DNA from TNF-α-induced (30 min) MEF nuclear extracts using RelA-specific antibody. The sequences DNA are indicated below. The high affinity κB-site highlighted in green, the low affinity κB-motif in blue, and mutations in red. **B.** Immunoblots from Flag pulldown assay with extracts of TNF-α-induced HEK293T co-transfected with Flag-tagged Nfatc1, Nfat5, Tead3, Cux1, or Smad4 and HA-tagged RelA. Transfection of empty vector was used as a control. **C.** Immunoblot analysis of endogenous RelA immunoprecipitated using anti-Flag beads from cell lysate of HEK293T cells transfected with Flag-tagged Nfatc1 or empty control vector. **D.** Pulldown assay using biotinylated wild-type Cxcl2 promoter DNA-streptavidin bead with nuclear extract of MEF cells without or with TNF-α stimulation. **E.** Native polyacrylamide EMSA with radiolabeled WT or double mutant Cxcl2 promoter DNA probes and purified recombinant p50:RelA heterodimer, Flag-NFATc1, or Flag-TEAD3. DNA sequences are listed in **A.** **F.** Native polyacrylamide EMSA with radiolabeled WT or mutant Cxcl2 promoter DNA (listed below) and recombinant p50:RelA and Flag-Nfatc1 (left) or Flag-Tead3 (right) * nonspecific protein band. DNA sequences are listed in **A**.

**Supplemental Figure 4.**
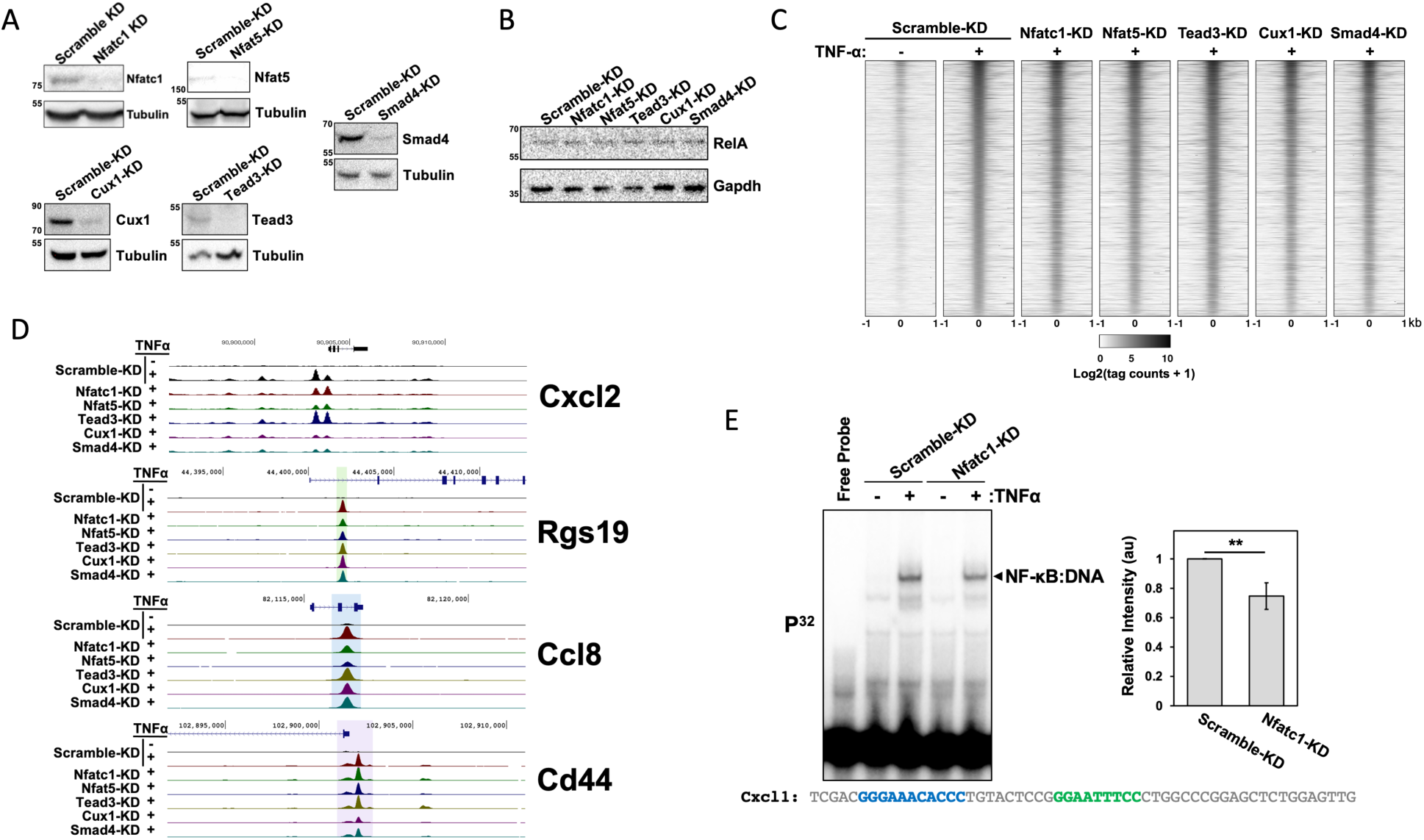
RelA ChIP of five cofactors and a scramble (control) KD MEF cell lines. **A.** Western blot of whole cell lysate from KD cell lines showing efficiency of KD. Level of tubulin is used to normalize loading. **B.** Western blot of whole cell lysate showing relative levels of RelA protein across all KD cell lines. Level of Gapdh is used to normalize loading. **C.** Histogram representation of log transformed RelA ChIP-Seq tag counts for RelA-induced peaks (log2FC > 1, p-value < 0.05, n = 14581 peaks in uninduced to TNF-α treated Scramble-KD cells) across all TNF-α stimulated KD cell lines. A window of +/- 1000 bp around the peak center is displayed. Peaks are sorted in decreasing order of total intensity observed in TNF-α stimulated control Scramble-KD cells. **D.** Genome browser tracks of RelA induced ChIP-Seq peaks in all KD cell lines showing differential effects of knockdowns at the target genes Cxcl2, Rgs19, Ccl8, and Cd44. RelA ChIP-Seq peak areas with differential reduction are highlighted in green (Nfat5-KD), blue (Nfatc1-KD and Nfat5-KD), and purple (Cux1-KD). **E.** (left) Native polyacrylamide EMSA with radiolabeled Cxcl1 promoter DNA (listed below) using nuclear extract of TNF-α stimulated (30 min) Scramble-KD and Nfatc1-KD MEF cell lines. (right) Quantification of relative NF-κB binding to Cxcl1 DNA with extracts of Scramble-KD and Nfac1-KD cell lines using a Bar Plot. **P < 0.01. P value was calculated by one-tailed Student’s t-test. N = 3 independent experimental replicates. Error bar represents standard deviation.

**Supplementary Figure 5.**
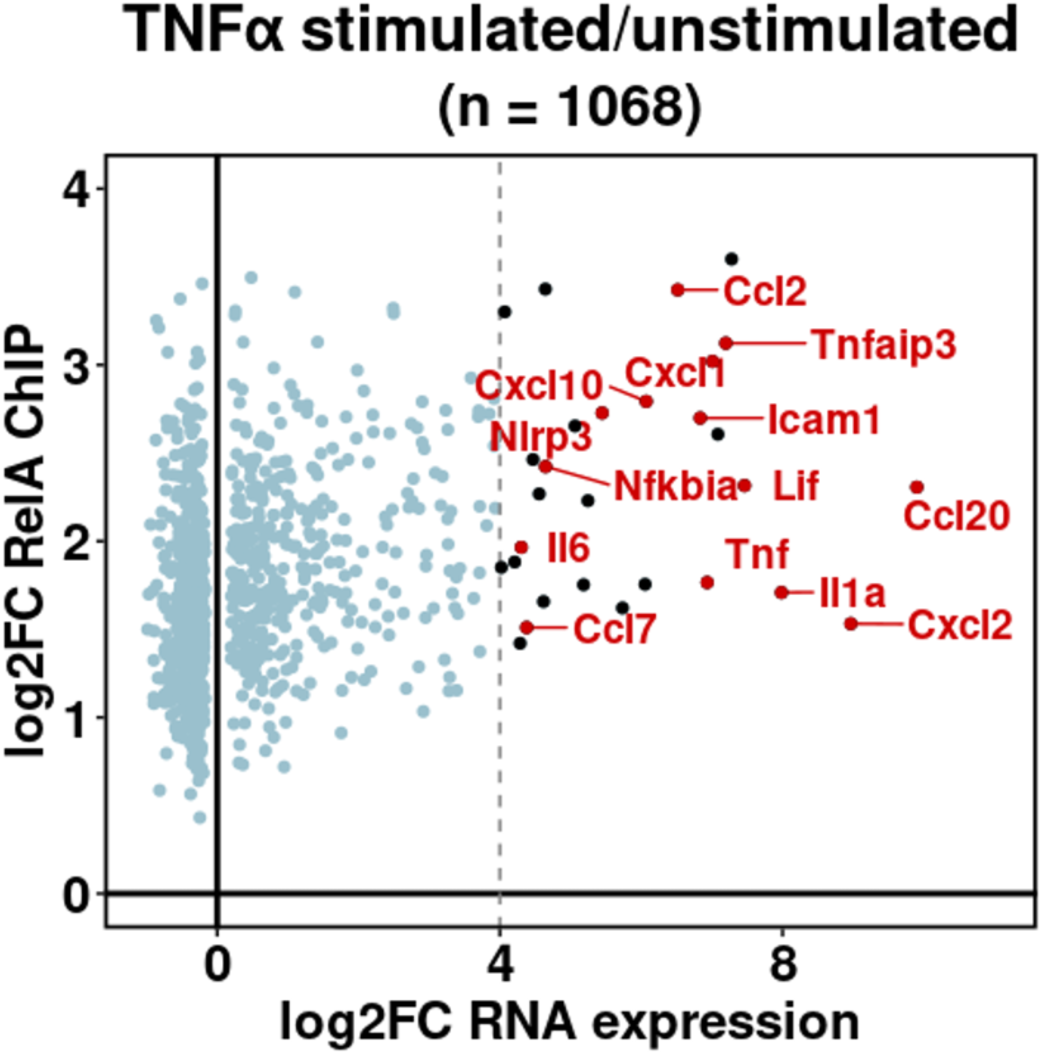
Correlation between changes in RelA ChIP-Seq signal and RNA expression in Scramble-KD cells. Genes were filtered to include significantly differentially expressed genes (p value < 0.05, Wald test) with induced RelA peaks (p value < 1E-5, Wald). Known NF-κB targets are labeled in red. Fold change calculations are based on triplicate RNA-Seq and duplicate ChIP-Seq datasets. n = 1068 data points analyzed.

**Supplementary Figure 6.**
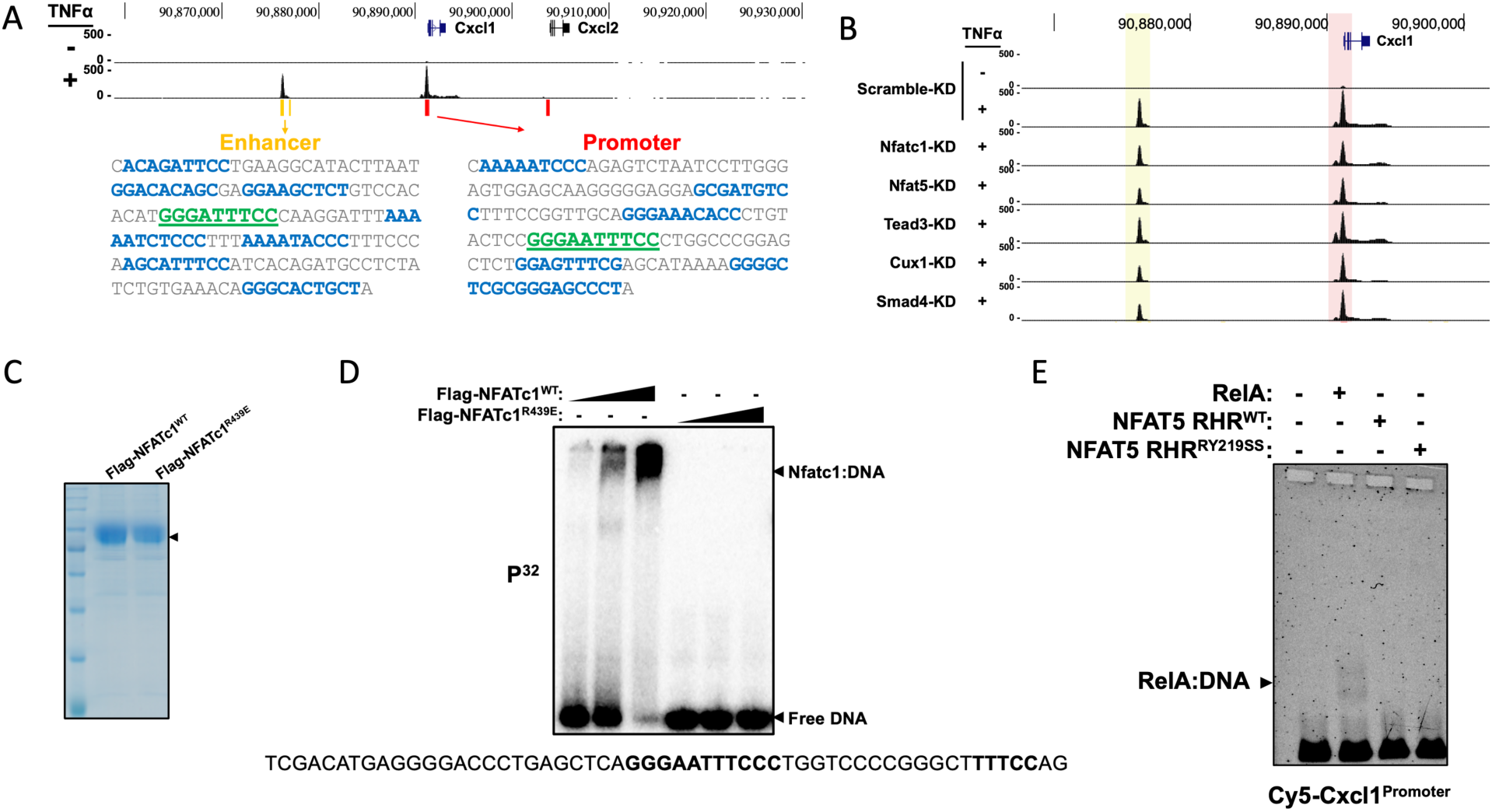
**A.** Genome browser representation of RelA ChIP-Seq signal at the Cxcl1-Cxcl2 locus in TNF-α stimulated (30 min) control cells. Yellow and red lines below tracks indicate predicted enhancer and promoter regions in the ENCODE database. The mm10 genomic sequences of the enhancer and promoter regions are also shown, with the central high-affinity κB motif highlighted in green and putative low-affinity κB sites in blue. The enhancer and promoter regions are separated by approximately 15 kbp. **B.** Genome browser tracks of RelA induced ChIP-Seq peaks at the Cxcl1 enhancer and promoter regions of all KD MEF cell lines. Red and yellow shades highlight RelA binding regions at the promoter and enhancer. **C.** Purified Flag-Nfatc1^WT^ and Flag-Nfatc1^R439E^ from transfected HEK 293T cells resolved in a 10% SDS-polyacrylamide gel and Coomassie stained. Cells were transfected for 24 h prior to harvesting for purification. The leftmost lane is of the molecular weight marker, indicating size of the purified proteins to be approximately 78 kDa. **D.** Native polyacrylamide EMSA of radiolabeled Cxcl1 promoter DNA (listed below) with increasing concentrations of purified Flag-Nfatc1, either WT or DNA-binding deficient mutant. The DNA sequence is displayed below with binding motifs for Nfatc1 in bold letters. **E.** Native agarose EMSA of Cy5-labeled Cxcl1 promoter DNA with purified recombinant His-tagged RelA, Nfat5 RHR, or DNA binding deficient mutant Nfat5 RHR^RY219SS^.

**Supplementary Table S3.** Sequences of DNA and DNA primers. (Table attached as a separate file denoted as SupplementaryTable3.xlsx)

**Supplementary Table S4.** GEO Series records for ChIP-seq Data: GSE287359 [ChIP-seq] Jan 22, 2025 approved XLSX

|  |  |  |  |  |
| --- | --- | --- | --- | --- |
| GSM8744856 | Scramble-KD MEF unstim ChIP-seq (replicate 1) | Jan 22, 2025 | approved | None |
| GSM8744857 | Scramble-KD MEF unstim ChIP-seq (replicate 2) | Jan 22, 2025 | approved | None |
| GSM8744858 | Scramble-KD MEF TNF ChIP-seq (replicate 1) | Jan 22, 2025 | approved | None |
| GSM8744859 | Scramble-KD MEF TNF ChIP-seq (replicate 2) | Jan 22, 2025 | approved | None |
| GSM8744860 | Nfatc1-KD MEF TNF ChIP-seq (replicate 1) | Jan 22, 2025 | approved | None |
| GSM8744861 | Nfatc1-KD MEF TNF ChIP-seq (replicate 2) | Jan 22, 2025 | approved | None |
| GSM8744862 | Nfat5-KD MEF TNF ChIP-seq (replicate 1) | Jan 22, 2025 | approved | None |
| GSM8744863 | Nfat5-KD MEF TNF ChIP-seq (replicate 2) | Jan 22, 2025 | approved | None |
| GSM8744864 | Tead3-KD MEF TNF ChIP-seq (replicate 1) | Jan 22, 2025 | approved | None |
| GSM8744865 | Tead3-KD MEF TNF ChIP-seq (replicate 2) | Jan 22, 2025 | approved | None |
| GSM8744866 | Cux1-KD MEF TNF ChIP-seq (replicate 1) | Jan 22, 2025 | approved | None |
| GSM8744867 | Cux1-KD MEF TNF ChIP-seq (replicate 2) | Jan 22, 2025 | approved | None |
| GSM8744868 | Smad4-KD MEF TNF ChIP-seq (replicate 1) | Jan 22, 2025 | approved | None |
| GSM8744869 | Smad4-KD MEF TNF ChIP-seq (replicate 2) | Jan 22, 2025 | approved | None |

**Supplementary Table S5.** GEO Series records for RNA-seq Data: GSE287360 [RNA-seq] Jan 22, 2025 approved XLSX

|  |  |  |  |  |
| --- | --- | --- | --- | --- |
| GSM8744870 | Scramble-KD MEF unstim RNA-seq (replicate 1) | Jan 22, 2025 | approved | None |
| GSM8744871 | Scramble-KD MEF unstim RNA-seq (replicate 2) | Jan 22, 2025 | approved | None |
| GSM8744872 | Scramble-KD MEF unstim RNA-seq (replicate 3) | Jan 22, 2025 | approved | None |
| GSM8744873 | Scramble-KD MEF TNF RNA-seq (replicate 1) | Jan 22, 2025 | approved | None |
| GSM8744874 | Scramble-KD MEF TNF RNA-seq (replicate 2) | Jan 22, 2025 | approved | None |
| GSM8744875 | Scramble-KD MEF TNF RNA-seq (replicate 3) | Jan 22, 2025 | approved | None |
| GSM8744876 | Nfatc1-KD MEF unstim RNA-seq (replicate 1) | Jan 22, 2025 | approved | None |
| GSM8744877 | Nfatc1-KD MEF unstim RNA-seq (replicate 2) | Jan 22, 2025 | approved | None |
| GSM8744878 | Nfatc1-KD MEF unstim RNA-seq (replicate 3) | Jan 22, 2025 | approved | None |
| GSM8744879 | Nfatc1-KD MEF TNF RNA-seq (replicate 1) | Jan 22, 2025 | approved | None |
| GSM8744880 | Nfatc1-KD MEF TNF RNA-seq (replicate 2) | Jan 22, 2025 | approved | None |
| GSM8744881 | Nfatc1-KD MEF TNF RNA-seq (replicate 3) | Jan 22, 2025 | approved | None |
| GSM8744882 | Nfat5-KD MEF unstim RNA-seq (replicate 1) | Jan 22, 2025 | approved | None |
| GSM8744883 | Nfat5-KD MEF unstim RNA-seq (replicate 2) | Jan 22, 2025 | approved | None |
| GSM8744884 | Nfat5-KD MEF unstim RNA-seq (replicate 3) | Jan 22, 2025 | approved | None |
| GSM8744885 | Nfat5-KD MEF TNF RNA-seq (replicate 1) | Jan 22, 2025 | approved | None |
| GSM8744886 | Nfat5-KD MEF TNF RNA-seq (replicate 2) | Jan 22, 2025 | approved | None |
| GSM8744887 | Nfat5-KD MEF TNF RNA-seq (replicate 3) | Jan 22, 2025 | approved | None |
| GSM8744888 | Tead3-KD MEF unstim RNA-seq (replicate 1) | Jan 22, 2025 | approved | None |
| GSM8744889 | Tead3-KD MEF unstim RNA-seq (replicate 2) | Jan 22, 2025 | approved | None |
| GSM8744890 | Tead3-KD MEF unstim RNA-seq (replicate 3) | Jan 22, 2025 | approved | None |
| GSM8744891 | Tead3-KD MEF TNF RNA-seq (replicate 1) | Jan 22, 2025 | approved | None |
| GSM8744892 | Tead3-KD MEF TNF RNA-seq (replicate 2) | Jan 22, 2025 | approved | None |
| GSM8744893 | Tead3-KD MEF TNF RNA-seq (replicate 3) | Jan 22, 2025 | approved | None |
| GSM8744894 | Cux1-KD MEF unstim RNA-seq (replicate 1) | Jan 22, 2025 | approved | None |
| GSM8744895 | Cux1-KD MEF unstim RNA-seq (replicate 2) | Jan 22, 2025 | approved | None |
| GSM8744896 | Cux1-KD MEF unstim RNA-seq (replicate 3) | Jan 22, 2025 | approved | None |
| GSM8744897 | Cux1-KD MEF TNF RNA-seq (replicate 1) | Jan 22, 2025 | approved | None |
| GSM8744898 | Cux1-KD MEF TNF RNA-seq (replicate 2) | Jan 22, 2025 | approved | None |
| GSM8744899 | Cux1-KD MEF TNF RNA-seq (replicate 3) | Jan 22, 2025 | approved | None |
| GSM8744900 | Smad4-KD MEF unstim RNA-seq (replicate 1) | Jan 22, 2025 | approved | None |
| GSM8744901 | Smad4-KD MEF unstim RNA-seq (replicate 2) | Jan 22, 2025 | approved | None |
| GSM8744902 | Smad4-KD MEF unstim RNA-seq (replicate 3) | Jan 22, 2025 | approved | None |
| GSM8744903 | Smad4-KD MEF TNF RNA-seq (replicate 1) | Jan 22, 2025 | approved | None |
| GSM8744904 | Smad4-KD MEF TNF RNA-seq (replicate 2) | Jan 22, 2025 | approved | None |
| GSM8744905 | Smad4-KD MEF TNF RNA-seq (replicate 3) | Jan 22, 2025 | approved | None |

